# Re-programming of *Pseudomonas syringae* pv. *actinidiae* gene expression during early stages of infection of kiwifruit

**DOI:** 10.1101/340018

**Authors:** Peter A. McAtee, Lara Brian, Ben Curran, Otto van der Linden, Niels J. Nieuwenhuizen, Xiuyin Chen, Rebecca Henry-Kirk, Erin A. Stroud, Simona Nardozza, Jay Jayaraman, Erik H. A. Rikkerink, Cris G. Print, Andrew C. Allan, Matthew D. Templeton

## Abstract

**Background:** *Pseudomonas syringae* is a widespread bacterial species complex that includes a number of significant plant pathogens. Amongst these, *P. syringae* pv. *actinidiae* (*Psa*) initiated a worldwide pandemic in 2008 on cultivars of *Actinidia chinensis* var. *chinensis*. To gain information about the expression of genes involved in pathogenicity we have carried out transcriptome analysis of *Psa* during the early stages of kiwifruit infection.

**Results:** Gene expression in *Psa* was investigated during the first five days after infection of kiwifruit plantlets, using RNA-seq. Principal component and heatmap analyses showed distinct phases of gene expression during the time course of infection. The first phase was an immediate transient peak of induction around three hours post inoculation (HPI) that included genes that code for a Type VI Secretion System and nutrient acquisition (particularly phosphate). This was followed by a significant commitment, between 3 and 24 HPI, to the induction of genes encoding the Type III Secretion System (T3SS) and Type III Secreted Effectors (T3SE). Expression of these genes collectively accounted for 6.3% of the bacterial transcriptome at this stage. There was considerable variation in the expression levels of individual T3SEs but all followed the same temporal expression pattern, with the exception of HopAS1, which peaked later in expression at 48 HPI. As infection progressed over the time course of five days, there was an increase in the expression of genes with roles in sugar, amino acid and sulfur transport and the production of alginate and colanic acid. These are both polymers that are major constituents of extracellular polysaccharide substances (EPS) and are involved in biofilm production. Reverse transcription-quantitative PCR (RT-qPCR) on an independent infection time course experiment showed that the expression profile of selected bacterial genes at each infection phase correlated well with the RNA-seq data.

**Conclusions:** The results from this study indicate that there is a complex remodeling of the transcriptome during the early stages of infection, with at least three distinct phases of coordinated gene expression. These include genes induced during the immediate contact with the host, those involved in the initiation of infection, and finally those responsible for nutrient acquisition.

## Background

*Pseudomonas syringae* is a widespread bacterial species complex that comprises plant epiphytes and pathogens, as well as being found in non-plant environments such as waterways [1, 2]. Each pathovar of *P. syringae* has a relatively narrow host range related to the specific effector and secondary metabolite profile encoded by its accessory genome. Effectors are proteins that are secreted into plant cells via the Type III Secretion system (T3SS) that function to repress the host defense response [3]. The kiwifruit vine (*Actinidia* Lindl spp.) disease pathogen *P. syringae* pv. *actinidiae* (*Psa*) was first identified in Japan in 1984 [4, 5] and was subsequently found in Korea in the 1990s [6]. Both these strains caused canker symptoms, but did not spread from their country of origin. In 2008, a particularly virulent canker-causing strain of *Psa* was reported in Italy and it quickly decimated plantings of *A. chinensis* var. *chinensis* cultivars, particularly ‘Hort16A’, ‘Hongyang’ and ‘Jin Tao’ [7]. This strain was found in other kiwifruit-growing regions including New Zealand, Chile and China by 2010 [8].

Whole genome sequence analysis was carried out on over 25 strains of *Psa* representing isolates from all locations where *Psa* had been reported. Phylogenetic analysis of the core genome indicated that the canker-causing isolates formed three clades. The first clade comprised the initial isolates from Japan, the second those collected in the 1990s from Korea and the third the pandemic outbreak strains from Italy, New Zealand, Chile and China [9–11]. Isolates within these clades are designated as biovars [12]. The core genome of isolates from the pandemic clade (biovar 3) differed by very few Single Nucleotide Polymorphisms (SNPs) suggesting that this is a clonal population; however, the isolates from New Zealand, Italy, Chile and China each possessed a different member of a family of integrative conjugative elements [9–11]. More recently a comprehensive phylogenetic analysis of eighty *Psa* isolates has shown the origin of the pandemic strains to be China [13]. Two new biovars of *Psa* have been recently discovered in Japan [14, 15], and thus the location of the source population of *Psa* biovars has yet to be conclusively determined.

The three canker-causing biovars each had a surprisingly varied accessory genome with different complements of genes encoding effectors and toxins [11, 16]. Many of these genes are encoded on putative mobile genetic elements. While bioinformatic analysis has identified genes that might be unique to the recent outbreak clade, little is known about the expression of these and other genes that might have a crucial role in pathogenicity. Surprisingly there are few RNA-seq data on the early stages of infection of plants by pathogenic bacteria, including *P. syringae*.

Several transcriptome studies have been carried out on different *P. syringae* pathovars [17, 18]. The most comprehensive *in planta* analysis has been of *P. syringae* pv. *syringae* (*Pss*) B728a. This pathovar is a particularly successful epiphyte as well as a pathogen of bean (*Phaseolus vulgaris* L.). Global analysis of the transcriptome as an epiphyte, pathogen, and under various stress conditions was carried out using a microarray covering >5000 coding sequences [19, 20]. The transcript profiles indicated that success as an epiphyte is enabled by flagellar and swarming motility based on surfactant production, chemosensing, and chemotaxis. This could indicate active relocation primarily on the leaf surface. Occupation of an epiphytic niche was accompanied by high transcript levels for phenylalanine degradation, which may help to counteract phenylpropanoid-based plant defenses [19]. In contrast, intercellular or apoplastic colonization led to the high-level expression of genes for *γ*-aminobutyric acid (GABA) metabolism (degradation of GABA would attenuate GABA repression of virulence) and the synthesis of phytotoxins, syringolin A and two additional secondary metabolites. Perhaps surprisingly the T3SS and T3SEs were not found to be strongly induced in the apoplast [19]. Subsequent analysis of several regulatory mutants illustrated a central role for GacS, SalA, RpoN, and AlgU in global regulation in *Pss* B728a *in planta* and a high degree of plasticity in these transcriptional regulators’ responses to distinct environmental signals [20].

More recently a comprehensive analysis of gene expression by *P. syringae* pv. *tomato* DC3000 (*Pto*) has been carried out on wild-type Arabidopsis and several defense gene mutants [21]. T3SS and T3SEs genes were upregulated *in planta*, as were transporter genes. A key finding was that Arabidopsis perturbs iron homeostasis in *Pto* [21].

To gain additional information about the expression of genes involved in pathogenicity, we have carried out transcriptome analysis of *Psa* grown *in vitro* on minimal media and *in planta* during the early stages of kiwifruit infection, using an RNA-seq approach. Our analysis showed that there are at least three distinct coordinated phases of gene expression and has resulted in the discovery of several uncharacterized genes that may have a role in pathogenicity.

## Results

### Infection assay and RNA-seq time course

Infection assays of kiwifruit tissue-cultured plantlets with *Psa* were performed in flood-inoculated tissue culture vessels under sterile conditions. This method gave consistent and reproducible infection rates, with water-soaked lesions on the underside of leaves progressively developing from day 5, necrotic lesions appearing from day 10 and plant death from four weeks (Additional file 1). A time course was carried out to assess the rate of infection and to measure and distinguish the relative populations of apoplastic and leaf surface-colonizing (epiphytic) bacteria. Bacterial counts in the apoplast (surface sterilized samples) rose rapidly during the first six days post inoculation and reached a plateau at approximately 10^8^ colony-forming units (CFU)/cm^2^ thereafter (Figure 1). Total bacterial counts (non-surface sterilized samples), which included both apoplastic and epiphytic bacteria, rose from 10^5^ to 10^8^ CFU/cm^2^ during the time course. The results suggest that for the first two days of infection, the majority of live cells were located epiphytically on the surface of the plant, but that the proportion of apoplastic colonizing bacteria progressively rose from days 2 to 6 so that the apoplast became the predominant niche from that time on.

**Figure 1.**
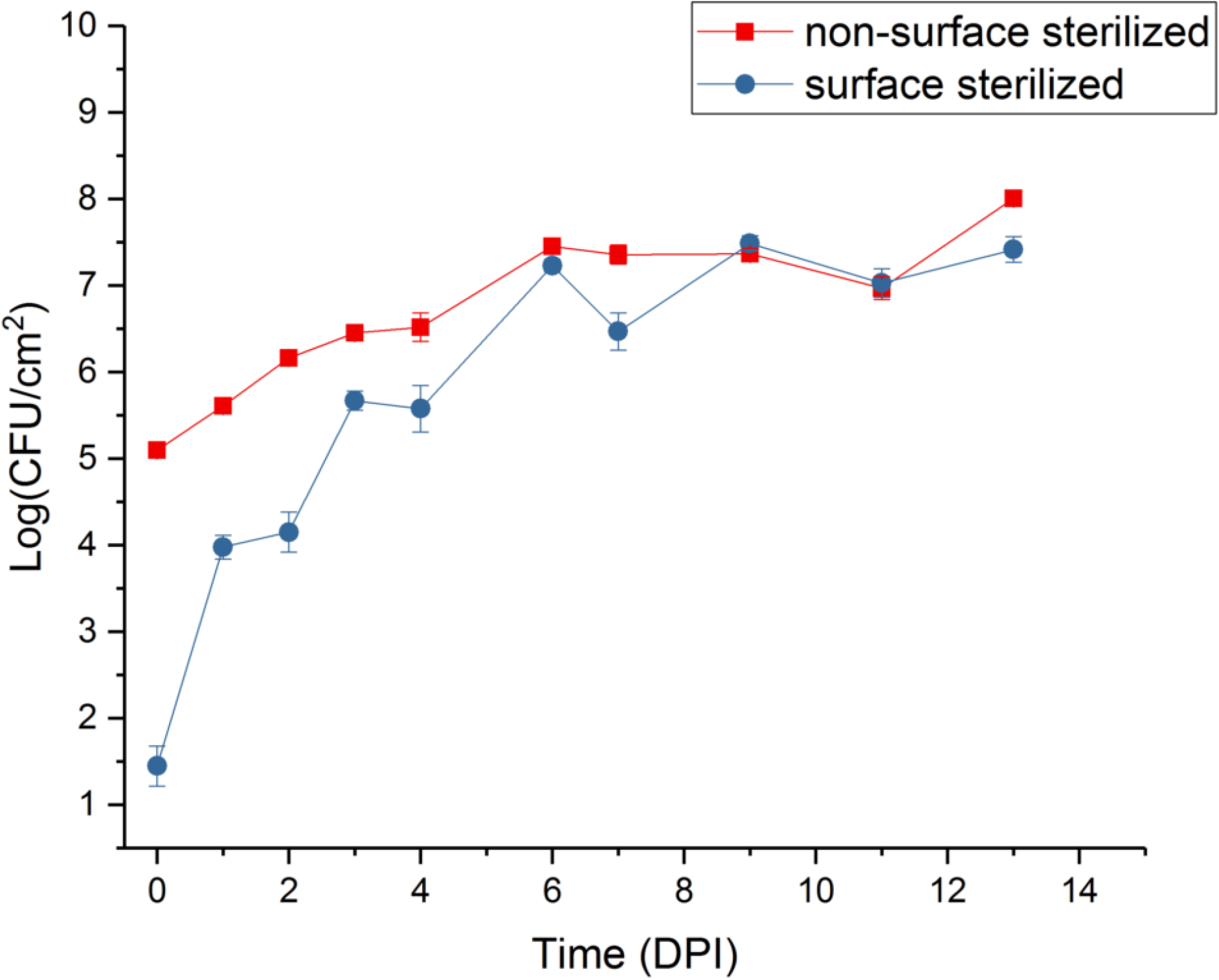
Time course of *Pseudomonas syringae* pv. *actinidiae* (*Psa*) infection of kiwifruit plantlets over 14 days. CFU, colony forming units; DPI, days post inoculation. 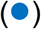 surface-sterilized; 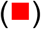 non-surface sterilized. The experiment was duplicated and each had four technical replicates. Error bars represent SE, n=8. Zero-time controls had no *Psa* present.

A time course of five days (120 hours) was selected for the RNA-seq analysis, with a focus on very early time points to identify genes induced in the first stages of contact with the plant surface and subsequent infection. It was postulated that key genes responsible for the initiation of infection would be induced at the early stages of contact with the plant surface. Obtaining significant numbers of bacterial reads from infected plants at the early stages of infection is extremely challenging. For this reason, leaves were not surface sterilized before RNA extraction.

### RNA-seq expression profile

Trimmed reads were mapped onto the complete *Psa* ICMP 18884 genome (CP011972.2 and CP011973) [22]. An average of 50,000-200,000 reads mapped to the *Psa* genome for each time point. This represents between 0.2 and 0.6% of the total reads per sample. The 27 control uninfected treatments showed 50-350 reads mapping to the *Psa* genome (0.0012-0.0002% of the total reads). A principal component analysis (PCA) was carried out on the inoculated samples to assess overall similarity, and the three biological replicates showed little variance within each time point (Figure 2A). PCA also demonstrated that each of the *Psa*-infected tissue samples belonged to one of three major phases that closely aligned to the post inoculation period and that were distinct from the *in vitro* control. The component groupings included the *in vitro* control, an early phase of infection (1.5 and 3 hours post infection, HPI), a mid-phase of infection (6, 12, and 24 HPI), and a late phase of infection (48, 72, 96, and 120 HPI) (Figure 2B).

**Figure 2.**
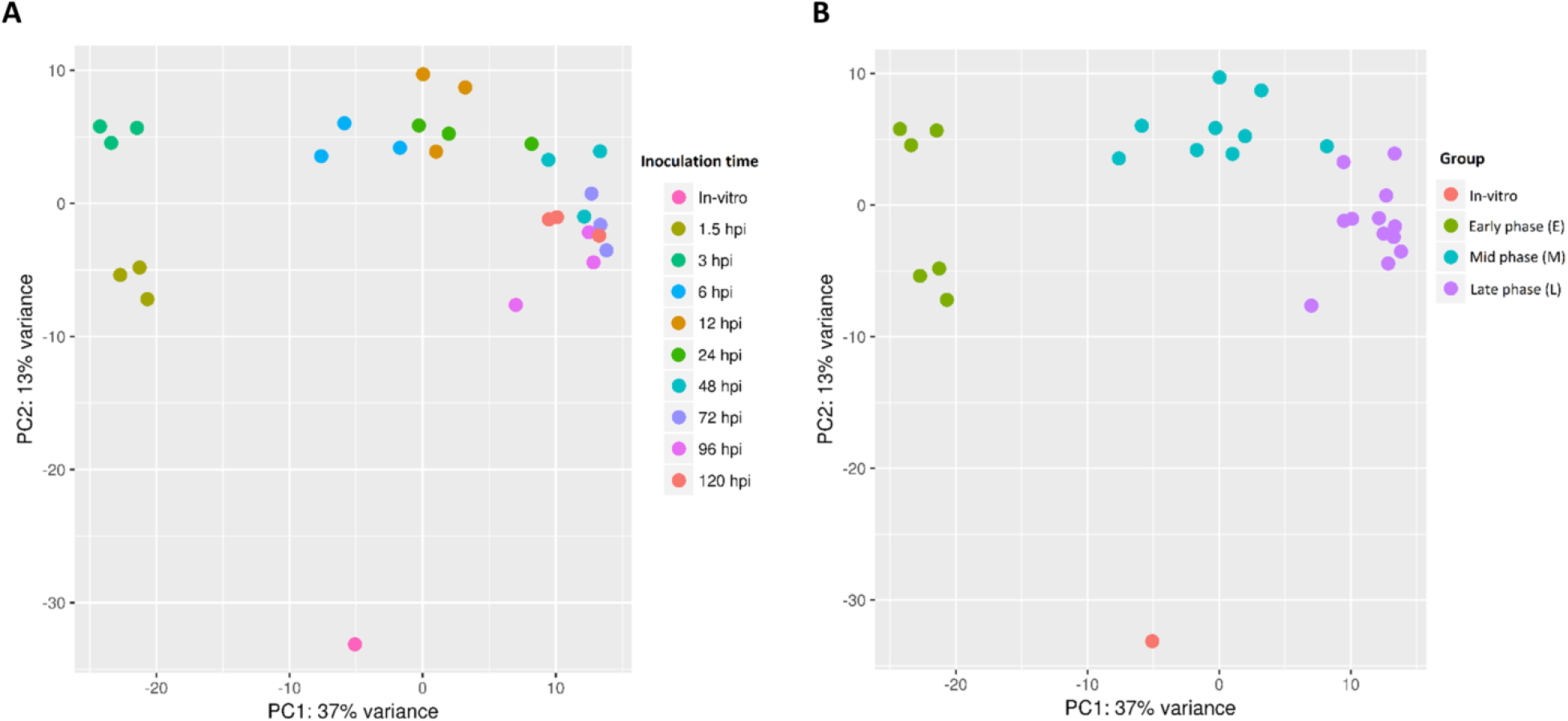
Principal component analysis plot (PCA) showing the clustering of VST (variance stabilizing transformation) transcriptomic data. (A) data points are colored by treatment time point (1.5 HPI, 3 HPI, 6 HPI, 12 HPI, 24 HPI, 48 HPI, 72 HPI, 96 HPI and 120 HPI). (B) data points are colored by infection phase (*in vitro*, Early (1.5-3 HPI), Mid (6-24 HPI), Late (48-120 HPI)). HPI, hours post infection.

### Heat map analysis with k-means clustering

Comparison of expression profiles is a powerful tool that can be used to identify and discover genes under the same regulatory regime. Furthermore, it was postulated that novel genes that showed similar expression profiles to known genes involved in pathogenicity might also have a role in causing disease. To identify such genes, similarities in the expression values for each gene were determined by first normalizing expression against its maximum value and then clustering by k-means analysis [23]. This analysis was restricted to those genes displaying Reads Per Kilobase per Million (RPKM) values over 50 for at least one time-point, to eliminate lowly expressed genes from the analysis [24]. Of the 5985 predicted gene models in the core and accessory genomes of *Psa*, 269 genes did not display evidence of being expressed at any sample point, 1473 had no sample point with an RPKM above 50, and 4243 genes had at least one sample point with an RPKM value above 50 (Additional file 2). Hierarchical Clustering on Principal Components (HCPC) using the remaining 4243 genes was used to partition a k-means analysis of genes into 13 clades (clusters) based on their expression profiles (Figure 3). These were further consolidated into six groups based on their broader expression patterns (Table 1). Of these 4243 genes, 1137 were constitutively expressed, 1323 genes were down-regulated *in planta*, and a further 815 did not show significant differential expression (Table 1). The remaining 968 genes were up-regulated *in planta* compared with *in vitro* and thus likely to have the most direct relevance to pathogenicity. Of the upregulated genes, there were three distinct groupings that differed in their temporal patterns and level of gene expression over the time course. These groups corresponded to 107 genes induced in the early (1.5 and 3 HPI) time points (early phase), followed by a group of 311 genes highly induced between 3 and 24 HPI (mid phase). The latter group included the majority of the T3SS and T3SE genes controlled by the HrpL regulon. Finally, 550 genes increased in their expression towards the late (48-120 HPI) time points (late phase). These three phases of gene expression were similar to the groupings identified by PCA analysis (Figure 2). Expression profiles were subsequently evaluated in more detail for genes with known or as yet undetermined roles in pathogenicity.

**Table 1.** k-means clustering of 4243 genes Psa into 6 groups. HPI, hours post infection.

| Group | k-means clade | Number of genes | Description | Expression Phase |
| --- | --- | --- | --- | --- |
| 1 | 1 & 13 | 1137 | Constitutively expressed genes | N/A |
| 2 | 2,3, & 4 | 1323 | Genes down-regulated *in planta* | N/A |
| 3 | 5 | 815 | Little differential expression compared to *in vitro* | N/A |
| 4 | 6, 10 & 11 | 311 | Genes upregulated (3-24 HPI) *in planta* | Mid |
| 5 | 7,8, & 9 | 550 | Genes upregulated late (48-120 HPI) *in planta* | Late |
| 6 | 12 | 107 | Early upregulated (1.5-3 HPI) genes *in planta* | Early |

**Figure 3.**
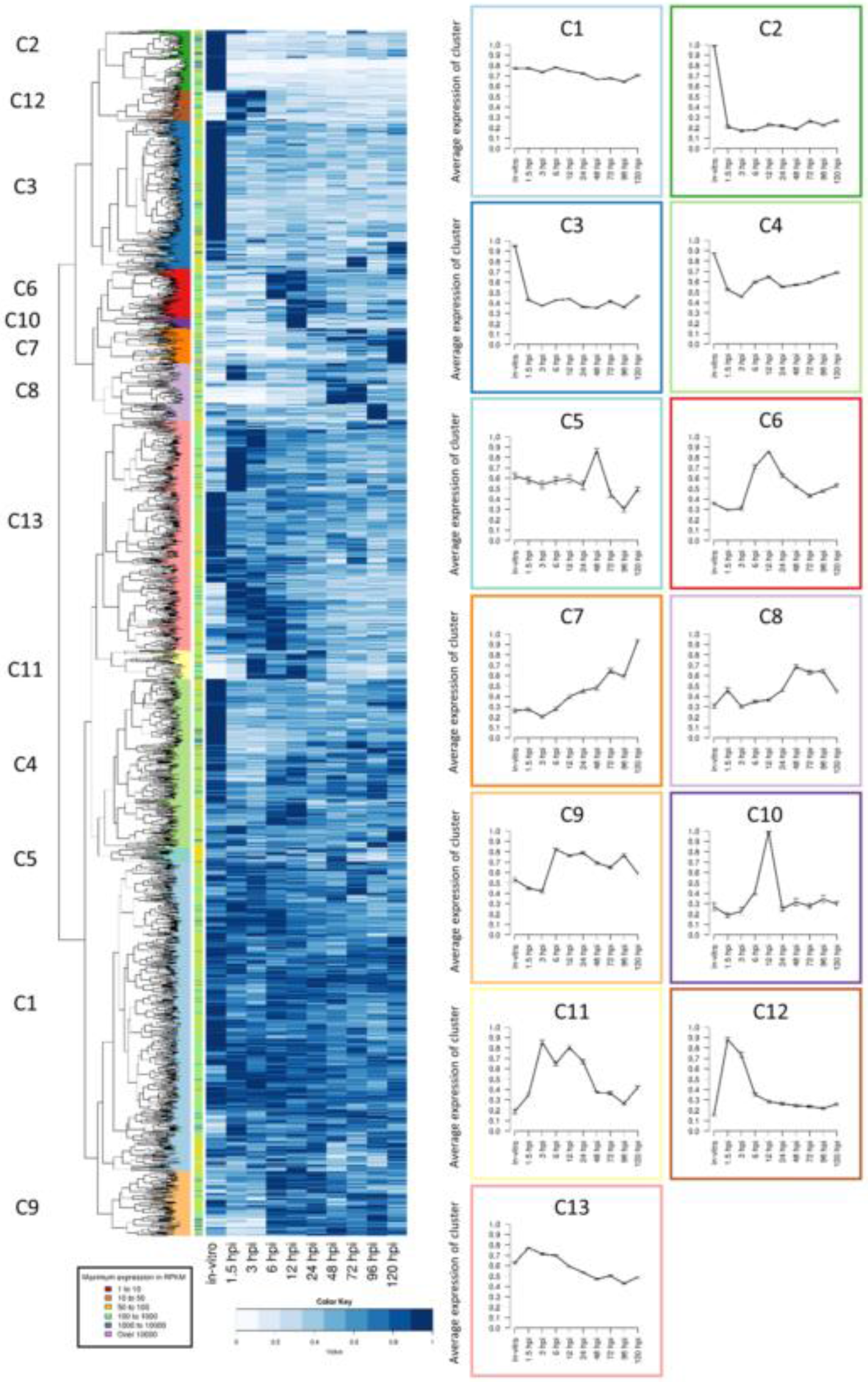
Heatmap and k-means clustering showing the expression of *Pseudomonas syringae* pv. *actinidiae* (*Psa*) genes in ‘Hort16A’ kiwifruit plantlets post infection. Similar expression profiles were clustered into 13 distinct groups by k-means. Line graphs displaying the prototype mean expression of each cluster (C) are included on the right. Error bars represent standard error.

### Early phase of infection characterized by the induction of a type VI secretion system and nutrient adaptation

Approximately 100 genes were up-regulated immediately upon contact with the host in the early phase of infection (1.5-3.0 HPI). These were found in clade 12 from the clustering analysis (Figure 3, Table 1). The majority of these genes were annotated as being involved in nutrient acquisition (Additional file 3). This probably reflects the adaptation to the surface of a leaf, where nutrients are scarce [25]. Genes that were particularly highly expressed included those predicted to be involved in phosphate and iron transport. In addition, some genes involved in the degradation of cell wall polymers, including a polygalacturonase (IYO_008325), were in this group. However, few genes predicted to have a direct role in pathogenicity were found. Two of the 43 annotated chemotactic response genes (chemoreceptors) found in the *Psa* genome were highly expressed during the early phase (Additional file 3), but these two are not amongst those previously functionally characterized [26]. These chemoreceptors could have a role in locating stomata or other potential sites of entry into the plant. Another set of genes that was highly induced in this phase encodes a putative Type VI Secretion System (T6SS). Effectors secreted through the T6SS have a variety of roles usually associated with killing both prokaryotic and eukaryotic cells [27]. Roles for T6SS effectors as virulence factors for animal pathogens have been well documented; however, there is as yet no evidence for an equivalent function in plant pathogens [28, 29]. Alternatively, the T6SS induced by *Psa* may have a role under field conditions in the antagonism of competing epiphytic microbes on the leaf surface.

### Mid-phase of infection characterized by expression of T3SS and T3SEs

Genes from three clades (6, 10 and 11) from the clustering analysis show increased expression 3-24 HPI (Table 1). Of these the most striking is a large transcriptional commitment to the induction of the T3SS apparatus and the expression of T3SEs, which were among the most highly upregulated genes within these time points (Figure 3; Additional file 4). Transcripts encoding for T3SS and T3SEs rose from 1.5 HPI, peaking between 3 and 12 HPI before falling to about half maximal levels for the remainder of the time course. Between 3 and 12 HPI these genes collectively accounted for 6.3% of the total reads (Additional file 4). Expression of *hrpA1* was by far the highest of all T3SS genes, accounting for over 50% of these reads. The HrpA protein comprises the needle of the T3SS apparatus. For plant pathogens the needle is much longer than that of animal pathogens because of the need to penetrate the host cell wall, thus presumably requiring higher expression of the corresponding gene [30].

The *Psa* biovar 3 genome has 40 genes encoding T3SEs, and 35 of these are predicted to encode full-length proteins [11], including one additional T3SE (*hopBN1*) recently identified (http://pseudomonas-syringae.org/). The expression profile of T3SEs during the mid-phase of infection followed that of the T3SS transcripts, rising rapidly after 1.5 HPI, with a maximum between 3 and 12 HPI, and then falling for the rest of the time course to around 20-40% of the highest level. The expression levels of each effector varied considerably: most were relatively abundant, in particular *hopAU1, hopS2, hopAO2, hopAZ1, hopZ5* and *hopF2, avrRmp1, avrB4* and *avrPto5* peaked at over 1000 RPKM (Figure 4, Additional file 5). However, several other effectors were weakly expressed during all the early phases of infection < 150 RPKM, such as *hopAH1* and *hopBB1-2* (Figure 4, Additional file 5). This may be due to lack of a role for these particular genes in the infection of kiwifruit, expression at the later stages of disease development (after 120 HPI), or a role in the infection of tissues other than leaves.

**Figure 4.**
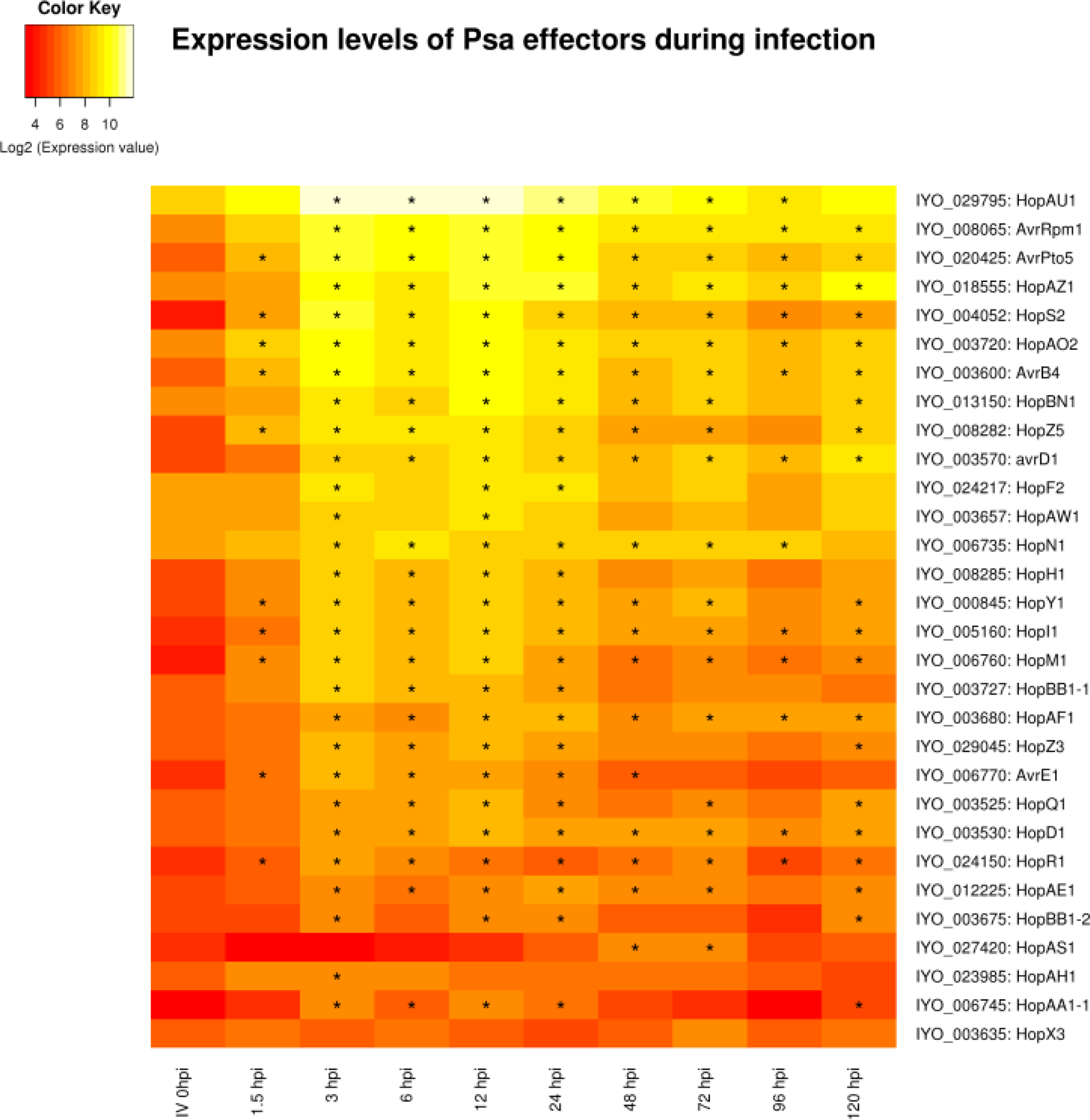
A heatmap showing the expression profiles of predicted *Pseudomonas syringae* pv. *actinidiae* (*Psa*) type III secretion system effectors at 10 time points during a infection time course of a Hotrt16a plantlets or in *in vitro* (IV)culture. This graph presents the log base 2 of the RPKM values with stars (*) indicating time points with significant (p < 0.01) changes in expression compared to the IV time point.

The effector that displayed the most distinct temporal expression profile was *hopAS1*. This full-length (1361 residue) effector had low expression in the second phase of infection and peaked at 48 HPI. *Pto* strains that are pathogenic on *Arabidopsis thaliana* carry a C-terminal truncated version of this effector (e.g. DC3000 402 residues) and hopAS1 is widely distributed in *P. syringae* [31]. Full-length versions caused effector-triggered immunity in almost all ecotypes of Arabidopsis, explaining why it effectively operates as a barrier to infection in this non-host. In contrast, deletion of the full-length version of *hopAS1* reduced virulence of *Pto* on tomato, suggesting it has a virulence function on this natural host. Both the *Pto* and *Psa* orthologs of this effector have a putative *hrp* box situated upstream of their putative start sites. In between lies a short uncharacterised potential open reading frame which could be an “unrecognised” effector chaperone. The *hopAS1* effector is also found in *P. syringae* pv. *phaseolicola*, where it was not found to be differentially expressed in response to induction of the HrpL TTSS regulatory system [32]. Unfortunately there are no strong clues about the possible biochemical function of *hopAS1*. It is one of the largest effectors (over 1300 residues, third largest *Psa* effector). Automated searching of the conserved domain database at NCBI identified just one tentative match (Bit score 51; E-value 5.7e^−6^) to a 330 residue portion of a heterodimerization domain in the N-terminus of the chromosome maintenance protein superfamily [33, 34]. Recently *hopAS1* was shown to be one of only six T3SEs from *Pto* able to bind to yeast plasma membrane, binding to several different phospho-inositol derivatives [35]. Unfortunately this research appears to have been performed with the truncated version of this gene from *Pto* DC3000. In contrast, the full-length *hopAS1* from *Psa* could not be localised when expressed in *Nicotiana benthamiana*, but this may be because it triggers cell death in that host [36].

T3SS and T3SEs are under the control of the HrpL regulon and hence their co-regulation would be expected. Several other genes that do not code for T3SEs also possess *hrp* boxes 5′ to their start site. These include genes that encode a putative lytic transglycosylase (IYO_006775), M20 peptidase (IYO_027210), *apbE* involved in thiamine biosynthesis (IYO_010630), a phosphatidylserine decarboxylase (IYO_025425), and an indole acetic acid-lysine ligase (*iaal*, IYO_002060, Additional file 6). *iaal* is found adjacent to a gene encoding a multidrug and toxic compound extrusion protein (*mate*, IYO_002055) on the chromosome of many *P. syringae* and *P. savastanoi* pathovars. Some *P. savastanoi* pathovars have an additional plasmid-associated *iaal* copy linked with indole acetic acid (IAA) production and gall formation. The proteins encoded by these genes are 92% identical, and the plasmid-located copy has been expressed heterologously and functionally characterized [37]. IAAL is postulated to convert free IAA into less active conjugate forms [38]. Heterologous expression of IAAL in tobacco and potato led to abnormal developmental changes [39]. Transcript levels of *Psa iaal* were induced early in infection and, in contrast to T3SS and T3SEs, remained high throughout the infection period; however, the adjacent *mate* gene did not appear to be highly expressed during this time period (Figure 5). In *Pto* DC3000 it has been shown that *iaal* can be both transcribed independently and co-transcribed with *mate* as an operon [40].

**Figure 5.**
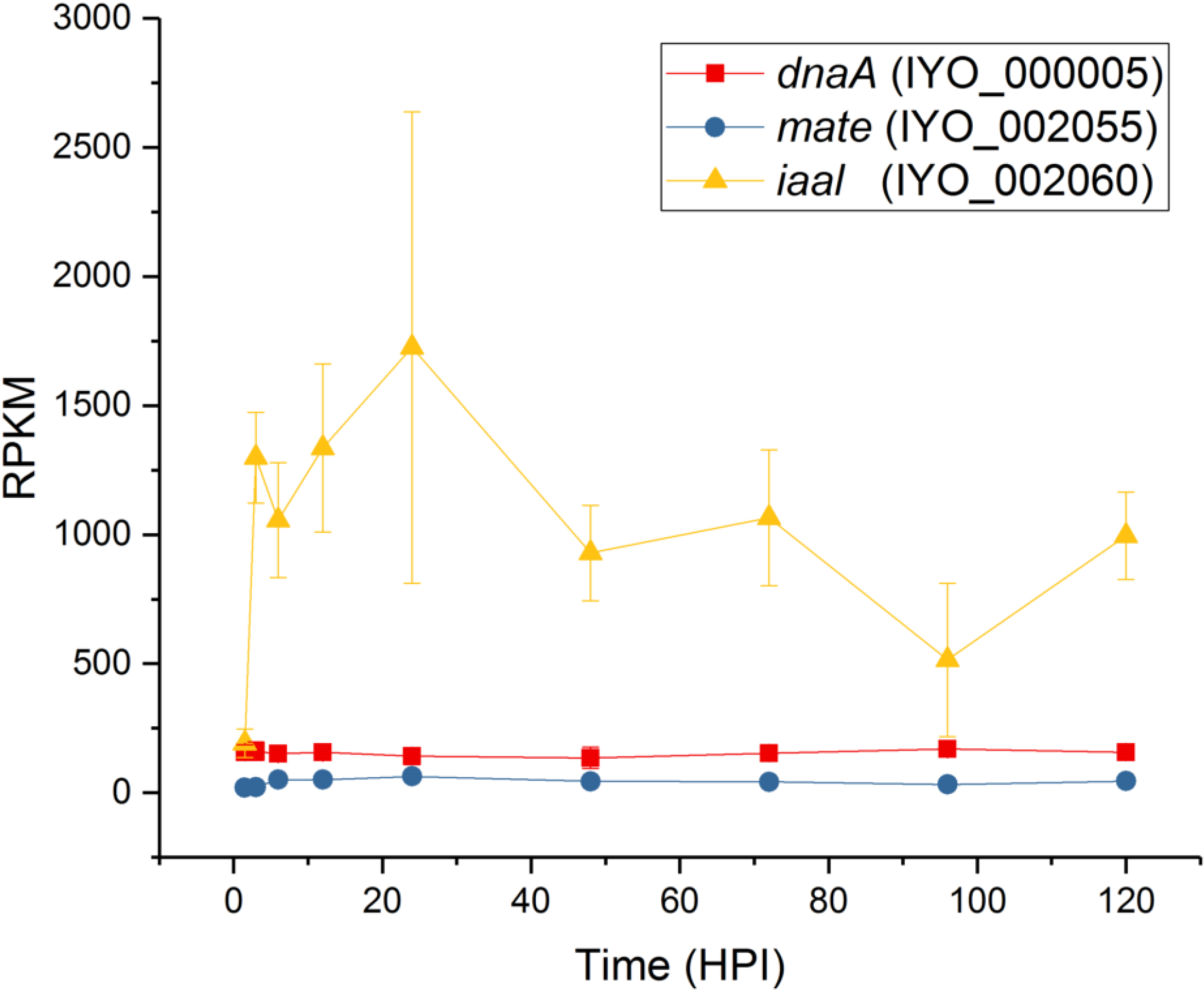
Expression of genes encoding *iaal* and *mate*. Reads per kb per million for *iaal* and *mate* were plotted over the infection time course. *dnaA* was included as a constitutively expressed control. Each point is the mean of three biological replicates with error bars representing standard error.

Other sets of genes that were strongly expressed during the mid-phase of infection (3-12 HPI) included four co-located genes on two operons that code for a diguanylate cyclase and two transcription factors, and thus may have a regulatory role (IYO_012110-25) (Additional file 7). Another set of four genes in two operons code for proteins involved in metal transport (IYO_003310-25); included in these is the highly expressed copper resistance/binding protein *copZ* (IYO_003325). Very high expression of the chemotaxis protein IYO_006420 was also observed; while not a membrane-bound chemoreceptor, it is predicted to contain a 4-helix bundle, which is a common chemoreceptor sensor domain [26]. This protein is predicted to be structurally similar to di-iron binding proteins (Pfam 09537), suggesting an alternate role in iron acquisition as opposed to chemoreception.

### Late phase of infection driven by nutrient acquisition and EPS production

A total of 550 genes were upregulated in the later phase of infection (groups 7-9). Ninety genes increased over 5-fold in expression between 1.5 and 120 HPI (Additional file 8). Of these genes, 14 were annotated to be involved in alginate and colanic acid biosynthesis and polymer export. Alginate is a hygroscopic polymer composed of D-mannuronic acid residues interspersed with L-guluronic acid residues with various degrees of acetylation [41]. This polymer has an important role in biofilm production and is well characterized in *P. aeruginosa* [42]. *P. syringae* is also known to produce alginate, but its role in pathogenicity is less well understood [43]. Recently it was shown **t**hat alginate accumulates in high amounts in the sub-stomatal spaces in *Psa-*infected leaves of kiwifruit (Sutherland et al., unpublished). A further 26 genes in this grouping were annotated as having a role in metabolite transport. This strongly suggests that as early-stage infection progresses there is a widespread induction of genes involved in metabolite transport and nutrient acquisition. These transporters are distinct from those observed in the early phase of leaf colonization.

### Expression of secondary metabolite pathways during infection

Several predicted secondary metabolite biosynthesis pathways have been identified in *Psa* using either antiSMASH 4.0 [44, 45], or by similarity to known biosynthetic pathways (Additional file 9). Three of these pathways, for achromobactin, pyoverdine and yersiniabactin, are involved in iron accumulation. *Psa* biovar 3 produces fluorescent compounds, i.e. pyoverdine, when grown on King’s B medium, but to a lesser extent than other *Psa* biovars [8]. The genes that code for this pathway appear to be poorly expressed *in planta* and on minimal media (Additional file 9). Genes coding for the alternative iron siderophores yersiniabactin and achromobactin are present in *Psa* but were also expressed at low levels *in planta* (Additional file 9). It has recently been postulated that plants are able to interfere with iron homeostasis in pathogenic bacteria, which may explain the apparent lack of expression of these pathways *in planta* [21]. Alternatively, *Psa* may be using a different mechanism for acquiring iron.

*Psa*-infected kiwifruit leaves show a distinct chlorotic halo which is presumably the result of the diffusion of a phytotoxin [8]. In addition to the three pathways with roles in iron absorption, there were four other secondary metabolite pathways identified in the *Psa* biovar 3 genome that have potential roles in pathogenicity and might account for leaf chlorosis. *Psa* biovar 3 possesses gene clusters involved in the biosynthesis of mangotoxin, a novel non-ribosomal peptide (NRP; IYO_003775-003830), an unknown metabolite (IYO_026725-026760), and an unknown compound synthesized from chorismate; the last-named pathway is plasmid-borne (Additional file 9). The genes involved in mangotoxin biosynthesis, NRP, and the unknown metabolite did not appear to be significantly induced during the early stages of infection, although genes in the NRP pathway were constitutively expressed between 50 and 100 RPKM throughout the infection time course (Additional file 9). While BLAST/antiSMASH searches did not identify likely products of either of the unknown biosynthetic pathways, many *Pseudomonas* spp. produce surfactive molecules to wet the leaf surface to aid motility [46]. In addition, the apoplast is a relatively dry environment that pathogens often modify to increase the relative humidity. For example, *syfA* - an NRP from *Pss* - produces an extremely hygroscopic molecule that facilitates wetting of surfaces including the leaf surface and apoplast [47–49].

The uncharacterized biosynthetic pathway on the plasmid of *Psa* biovar 3 has two operons, and is adjacent to a *luxR* receptor [50]. The first operon codes for a chorismate-utilizing enzyme and a glutamine amidotransferase (annotated as anthranilate synthase I and II) [11]. The second operon codes for the biosynthesis and secretion of a putative aromatic, but uncharacterized, compound that was strongly induced *in planta* after 12 HPI and remained steady for the remainder of the time course (Figure 6, Additional file 9). The plasmid-localized secondary metabolite pathway is not widespread in *P. syringae* but interestingly is also present in the vascular pathogen *Xylella fastidiosa*, the causal agent of Pierce’s disease, and some root-associated *Pseudomonas* species [11].

**Figure 6.**
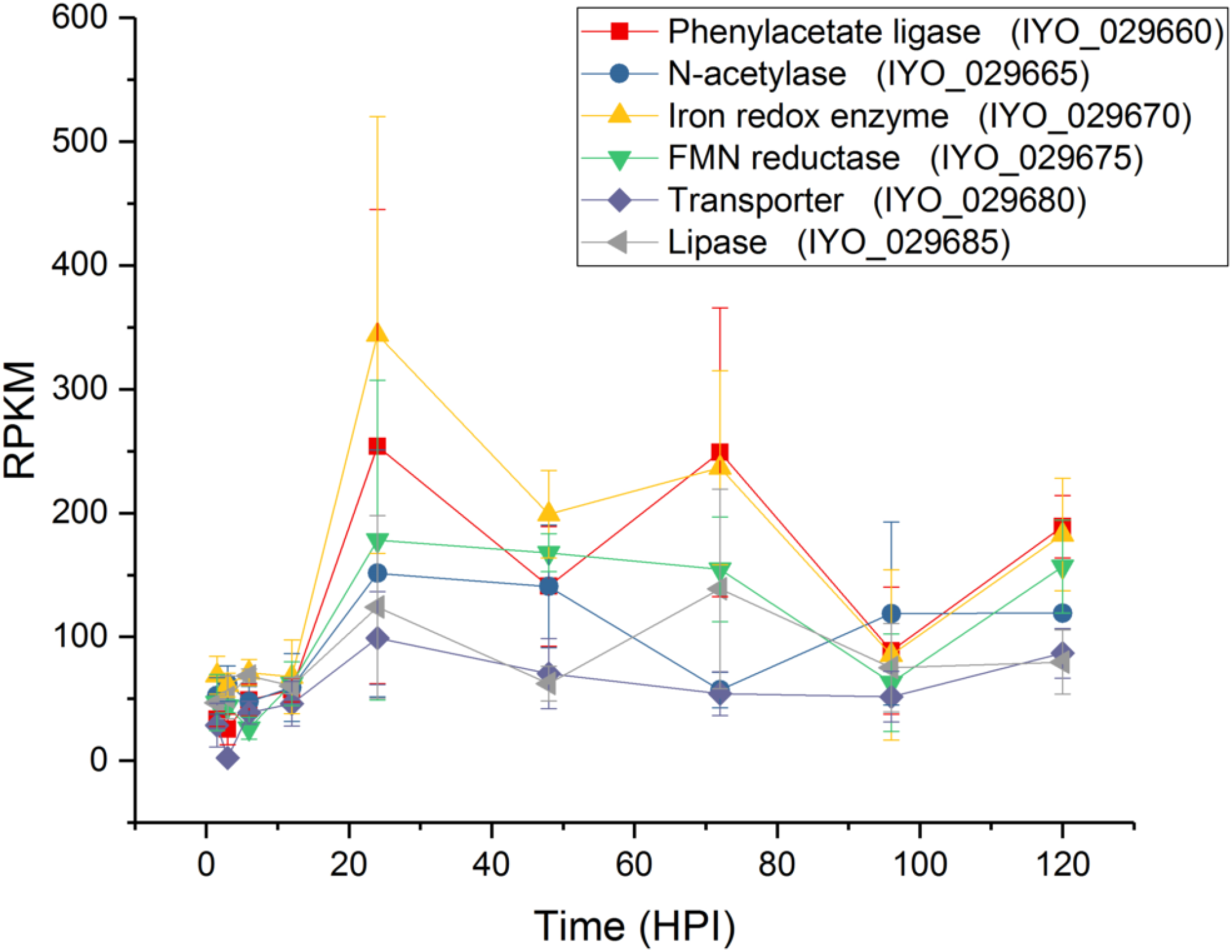
Expression of genes from the plasmid-borne putative aromatic biosynthetic pathway. Reads per kb per million values for the genes in the operon coding for the biosynthetic pathway of a putative aromatic compound plotted over the infection time course. Each point is the mean of three biological replicates with error bars representing standard error.

### Proteins secreted through the type II secretion system

In addition to the translocation of proteins through specialized structures such as the T3SS and the T6SS, bacteria also use the Sec or Tat systems to secrete proteins into the periplasm for the Type II secretion system (T2SS) to export [51]. This system will target proteins *in planta* to the apoplast, as opposed to the cytoplasmic location of the T3SEs. This is an important function since the plant apoplast has a number of largely constitutive antimicrobial defenses such as phytoanticipins, hydrolytic enzymes and enzyme inhibitors that may need to be inactivated to facilitate colonization. Recently analysis of the *Pto* secretome identified a protease inhibitor Cip1 as playing a role in virulence of *Pto* on tomato [52]. We were therefore interested to see if there were T2SS proteins upregulated in the early and mid-phases of infection.

Type II secreted proteins (T2SP) can be identified by their canonical secretory leader sequence using SignalP [53]. Five hundred and thirty-nine proteins were predicted to be secreted. Of these proteins, 21 were induced in the early phase of infection (clade12). Of the significantly induced genes (ratio RPKM 3HPI/RPKM *in vitro* >5), the majority are predicted subunits of membrane-bound complexes with a role in nutrient transport (Additional file 10). All of these had annotations assigned.

Twenty-six proteins with predicted leader sequences were present in the mid phase of infection (Additional file 10). Of those strongly expressed compared to *in vitro* growth, four were annotated as hypothetical proteins. However, two of these predicted gene products have similarity to enzyme inhibitors. IYO_001870 has homology to a superfamily of vertebrate lysozyme inhibitors and IYO_009660 contains a region with homology to Pfam domain 13670, present in some putative protease inhibitors. Both these proteins may have a role in neutralizing the apoplast and are candidates for further functional analysis. Forty-two non-annotated secreted proteins identified in the *Pto* genome were screened for the ability to inhibit the tomato C14 defense-related protease and one, Cip1, was shown to be an inhibitor [52]. Of these, 37 had orthologs (95% sequence identify) in *Psa* biovar 3 but only seven were clearly differentially expressed *in planta*. Interestingly, the *Psa* ortholog of Cip1 (IYO_021465) was not differentially expressed during the time course in this study.

### Validation of expression profiles using reverse transcription quantitative PCR

An independent infection time-course experiment was undertaken in order to validate the RNA-seq expression data. A total of 8 genes were selected for reverse transcription quantitative PCR (RT-qPCR) with their expression being assayed *in vitro* (IV) as well as at 2, 24 and 72 HPI (Figure 7, Additional file 13). These time points represented the early, mid and late phases of infection previously described. When compared to the normalized expression ratio of the RNA-seq data, the normalized qPCR data from 6 of these genes displayed a regression value above 0.7 indicating a high level of consensus between the RNA-seq and qPCR datasets for these genes. Two of the selected genes (IYO 000005 and IYO 003790) displayed regression values below 0.5, however the regression of IYO 000005 increased to greater than 0.7 if the outlying IV time point was omitted. These results show that the expression profile of the bacteria at each infection phase is largely consistent with a different assay methodology and independent experimentation.

**Figure 7.**
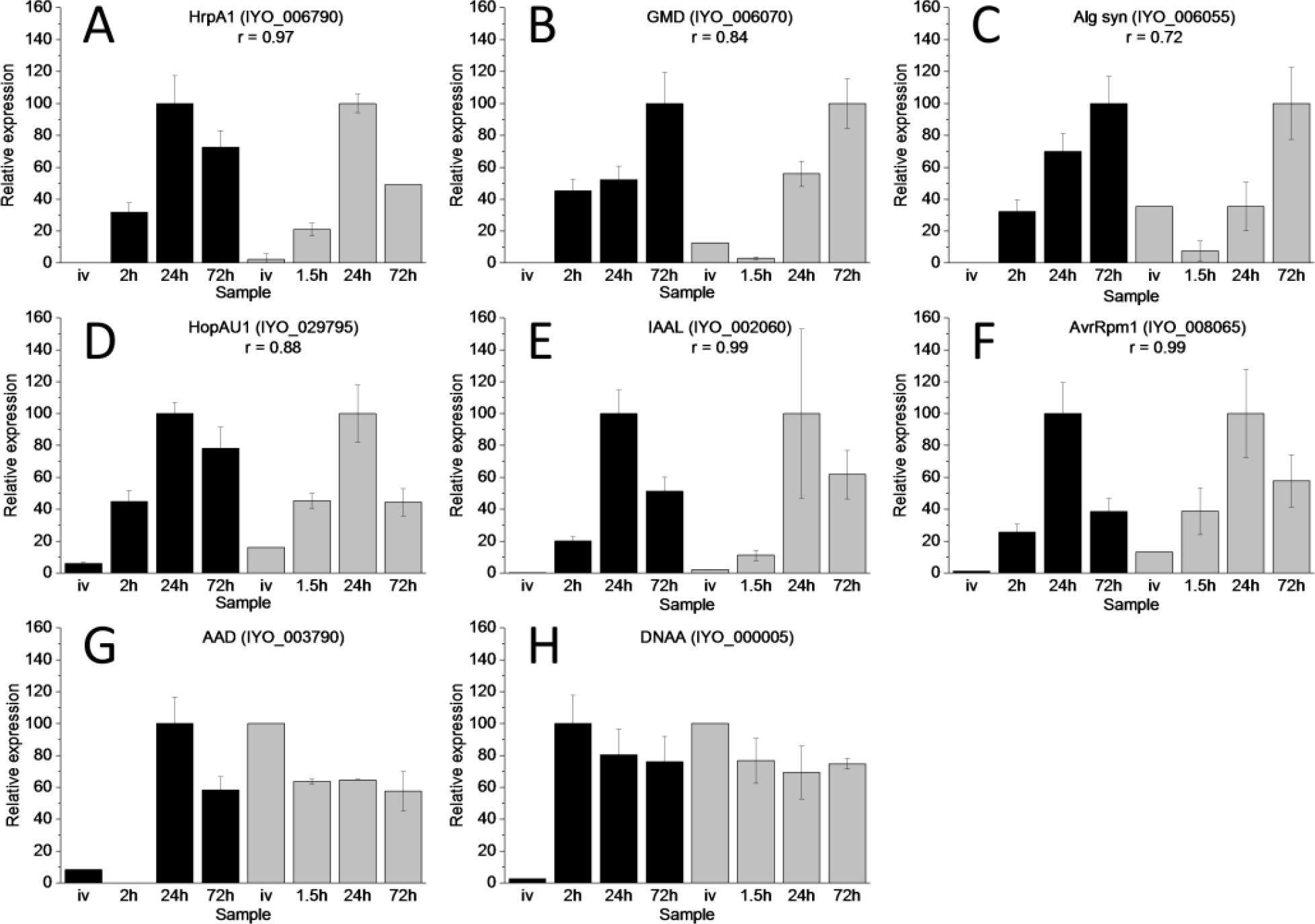
Reverse transcription quantitative PCR validation of RNA-seq data. Eight target genes and five potential reference genes were selected for reverse transcription quantitative PCR (RT-qPCR) (supplemental data set 13). RNA was extracted from *Psa* grown *in vitro* (IV) infected ‘Hort16A’ plantlets at 2, 24 and 72 hours post infection (HPI). RT-qPCR values (three biological replicates) were based on normalisation against the geometric average/mean of three reference genes (IYO_010670, IYO_009010 and IYO_002170) selected using Bestkeeper and geNorm analysis [69, 70]. For comparison with the RNA-seq data, values were displayed by representing maximum expression of each gene as 100. Pearson correlation coefficients (r) > 0.5 between the RT-qPCR and RNA-seq data are displayed all primers are listed in supplemental data set 13. RT-qPCR data is represented by black bars and RNA-seq (reads per kb per million) by grey bars. A: HrpA1 (IYO_006790); B: GDP-mannose dehydrogenase (GMD, IYO_006070); C: Alginate synthase (IYO_006055); D: HopAU1 (IYO_029795); E: IAAL (IYO_002060); F: AvrRpm1 (IYO_008065); G: Amino acid adenylation protein (AAD, IYO_003790); H: DNAA (IYO_000005).

## Discussion

RNA-seq was used to investigate the early stages of infection of kiwifruit plantlets by *Psa* biovar 3. This biovar is highly virulent on kiwifruit, with apoplastic CFU reaching a plateau from 6 days post inoculation in plantlet leaves. PCA and clustering analysis revealed three phases of gene expression *in planta* during early stages of colonization by *Psa*. The first was a rapid transient phase that occurred immediately upon contact with the plant. Included in these genes was a T6SS, which might have a role in pathogenesis, similarly to that in animal systems [27]. Alternatively, the T6SS may have a role in competition against epiphytic bacteria. Interestingly none of the other genes in this group had a predicted function that could have a direct role in pathogenicity. This suggests that these early expressed *Psa* genes play a role in rapid adaptation to the plant surface, since most of the bacterial counts were on the surface of the plant rather than the apoplast at this stage. Two chemotactic receptors were also highly expressed in this early phase. These are strong candidates for a role in sensing stomata, hydathodes and other points of entry for the pathogen.

In contrast, the mid phase of infection, which occurred between 3 and 12 HPI, included the T3SS and majority of the T3SEs. These genes were the most upregulated at these time points, accounting for over 6% of the transcripts detected. Similar results were observed for *Pto* colonization of Arabidopsis [21]. This is in contrast to the data observed for *Pss* during the early stages of infection of bean, where a large induction of either T3SS or T3SEs was not observed. This might be due to different infection strategies of the two pathogens: *Pss* is regarded as a stronger epiphyte than other *P. syringae* pathovars, because the phylogroup it belongs to (II) has a greater focus on toxin production, and its members typically have fewer effectors than other *P. syringae* phylogroups [1]. The difference could alternatively be attributed to the different experimental approaches employed [19]. Levels of individual *Psa* T3SE gene expression varied considerably during the time course. However, the temporal expression pattern was largely consistent between effectors, with most (25/30) fitting into the mid-phase gene expression clusters 6, 10 and 11. The most notable exception was *hopAS1*, which peaked later in expression around 48 HPI, as opposed to 3-12 HPI for most other effectors.

The roles of the T3SS and T3SE in repressing the induced host defense response are increasingly well understood. Less well understood is the repression of constitutive plant defenses in the apoplast, the inactivation of which is an essential prerequisite to the establishment of the T3SS. These defenses include phytoanticipins, cell wall degrading enzymes, proteases and enzyme inhibitors. Furthermore, the apoplast is a relatively dry space that needs to be humidified to optimize colonization [48, 49]. Two resident proteins in the conserved effector locus (CEL), *hopM* and *avrE*, appear to be important in establishing the right humidity conditions in other *P. syringae* hosts [54], and in *Psa* these effectors both follow the mid-peak expression profile but show only average expression levels. This study has also identified two predicted proteins that may have a role in neutralizing the apoplast. One was a predicted lysozyme inhibitor (IYO_001870) and the other a predicted protease inhibitor (IYO_009660). It is likely, however, that there are further genes to be discovered that play a role in neutralizing the apoplast, including the production of potential surfactants.

The final phase comprised of genes whose expression progressively increased over the five-day (120 h) time course. Included were a raft of genes coding for proteins involved in nutrient acquisition such as transporters. Notably, these were different transporters from those induced at the very early phase of infection. There was also strong induction of genes involved in alginate and colanic acid production. These compounds are a large component of the extracellular polysaccharide substances (EPS) and known virulence factors of *Pseudomonas* [41, 43]. Their precise role in pathogenicity is not known, but they have been postulated to protect the bacteria from adversity, in this case plant defenses, and also to enhance adhesion to solid surfaces. Indeed alginate synthesis, along with ice nucleation, auxin synthesis and auxin inactivation by IAAL, is common among the canonical *P. syringae* lineages that have been traced back to a last common ancestor (LCA) 150-180 million years ago [55]. Another component predicted to be derived from the LCA is the tripartite pathogenicity island structure consisting of the *hrp*/*hrc* gene cluster flanked by both the CEL and an exchangeable effector locus (EEL). *Psa* shows this tripartite structure, albeit that the EEL is further away from the other two pathogenicity islands. The heat map analysis did not highlight any obvious differences in expression patterns between the effectors located on these three pathogenicity islands.

While effectors are well known to play a role in plant defense suppression, the role of many other genes expressed during infection is far less certain. This study has identified a number of non-effector genes that were strongly induced *in planta* and are likely to be having a role in establishing infection. The relative importance of these will need to be ascertained using either gene knockouts or TraDIS (Transposon Directed Insertion-site Sequencing) [56, 57].

## Conclusions

The results from this study indicate that there is a complex remodeling of the bacterial transcriptome during the early stages of infection, with at least three distinct phases of coordinated gene expression. The first describes genes induced during the immediate contact with the host. These include the expression of a T6SS and genes annotated as involved in nutrient transport. The second phase was dominated by genes predicted to have roles in initiating infection and includes the T3SS and T3SEs. Included in this group are novel proteins that may have roles in neutralizing constitutive defenses in the apoplast. The final phase includes genes involved in nutrient transport and biofilm formation.

## Experimental Procedures

### Infection assays

*Actinidia chinensis* Planch. var. *chinensis* ‘Hort16A’ plantlets, grown from axillary buds on Murashige and Skoog rooting medium without antibiotics in a 400-mL clear plastic tub with a sealed lid, were purchased from Multiflora (http://www.multiflora.co.nz/home.htm). Plantlets were grown at 20°C under fluorescent lights with a 16 h on/8 h off regime and used within a month of purchase. For inoculation an overnight shake culture of *Psa* ICMP 18884 [11, 22] was grown in liquid Lysogeny Broth (LB) [58] at 20°C and 180 rpm shaking. The cell density was determined by measuring the absorbance at 600 nm. Cells were washed in 10 mM MgSO_4_ and resuspended at a cell density of 10^7^ CFU/mL (A_600_ 0.01). The surfactant Silwet L-77 (Cat VIS-30, Lehle Seeds, Round Rock, TX, USA) was added to the inoculum to a concentration of 0.0025% (v/v) to facilitate leaf wetting. The inoculation method was modified from that developed for Arabidopsis [59]. Containers with ‘Hort16A’ plantlets were filled with the inoculum fully submerging the plantlets and left for three minutes. Containers were drained, the lid replaced, then incubated in a controlled climate room at 20°C with a light/dark cycle of 16 h on/8 h off.

### Growth assay

Leaf samples were taken at different times post inoculation as appropriate. Each sample consisted of four leaf discs, taken with a 1-cm diameter cork borer, from four different leaves. All four discs were taken from the same tub. To estimate CFU, the apoplast discs were surface sterilized in 70% (v/v) ethanol for 30 s and subsequently washed in sterile Milli-Q water. Samples for estimation of total bacteria were not surface sterilized. Leaf discs were placed in Eppendorf tubes containing three stainless steel ball bearings and 300 μL 10 mM MgSO_4_, and macerated in a bead crusher for 2 min at maximum speed (Storm 24 Bullet Blender, Next Advance, Averill Park, NY, USA). A dilution series of the leaf homogenates was made in sterile 10 mM MgSO_4_ until a dilution of 10^−8^. The dilution series was plated in 5-μL droplets on LB medium supplemented with both 12.5 μg/mL nitrofurantoin and 40 μg/mL cephalexin. After 72 hours of incubation at 20°C CFU were counted for the lowest possible dilution(s), which was calculated back to the CFU per cm^2^ of leaf area.

### RNA extractions

RNA was extracted from *Psa* ICMP 18884 grown to late log phase at 18°C on Hoitnik and Sinden minimal media [60]. Cells were harvested and total RNA extracted using an Ambion RNA extraction kit (Thermo Fisher, Waltham, MA, USA). RNA was extracted from 1-month-old *A. chinensis* var. *chinensis* ‘Hort16A’ plantlets propagated from tissue culture infected with *Psa* as described above after 1.5, 3, 6, 12, 24, 48, 72, 96 and 120 HPI. Each time point consisted of three biological replicates. Three pots were used for each time point and each biological replicate consisted of three combined plantlets sampled across each of the three pots (Additional file 11). Mock-inoculated plants (submerged in 10mM MgSO_4_ only) were used as controls for each time point. RNA was extracted using the Spectrum™ Plant Total RNA Kit (Sigma-Aldrich, Milwaukee, WI, USA). Sequencing libraries were constructed from total RNA using the Ribo-Zero Plant procedure (Illumina, San Diego, CA, USA).

### Bioinformatics and differential expression analysis

Sequencing was performed using HiSeq2000 (Illumina) by Macrogen (www.macrogen.com). Raw RNA reads (100 bp paired end reads) were independently trimmed, quality filtered and had their adaptors removed using Trimmomatic v0.36 [61]. The trimming process involved removing the Truseq adaptor sequences and the first 15 base pairs of each read sequence. Regions of low quality calling were also removed from each read using a sliding window of one nucleotide and a quality score above 20. Reads with a length greater than 30 base pairs were selected for alignment following the trimming process.

Processed read sequences were aligned to the *Psa* ICMP 18884 genome sequence [22] using the Bowtie2 v2.25 aligner. HTSeq v0.9.1 was used in conjunction with the BAM outputs generated by Bowtie2 to count the number of alignments against the gene features defined in the corresponding *Psa* ICMP 18884 GFF3 file that had a mapping quality greater than 10 [62].

The count outputs for each gene feature acquired using HTSeq were used for differential expression analysis, principle component analysis, and for the calculation of Reads Per Kilobase per Million (RPKM) values. RPKM values were calculated by dividing the read alignment count for each gene feature in each library by the total number of reads in that library per million and then dividing this by the length of the gene in kilobases. Analysis of mapped read counts was done using the statistical software R (version 3.4.3). Principal component and differential expression analysis was done using the R package DESeq2 (v1.18.1) [63].

K-means cluster analysis of expression data was done using the R packages ggplot2 (v2.2.1) [64], FactoMineR (v1.39) [65] and FactoExtra (v1.05) [66]. K-means analysis used the average expression of each gene across biological replicates that was normalized against the time point that displayed the highest level of expression for that gene. Hierarchical clustering on principal components was used to decide the ‘optimum’ minimum number of clusters (k) to partition the output of k-means analysis into. This was achieved by calculating the sum of the within-cluster variance with increasing numbers of clusters. These within cluster variances were plotted as a bar plot (Additional file 12) in order to determine the cluster number(s) that produced a notable loss of inertia (variance). A notable loss of inertia was observed at a cluster number of 2, 3, 6, 7, and 13. Partitioning of the data using a cluster number less than 13 did not adequately describe the patterns of expression in enough detail, while greater numbers generated unnecessary complexity. It was hence decided that a cluster number of 13 represented the optimal minimum number of clusters.

### RT-q-PCR validation of RNA-seq data

To validate the RNA-seq dataset, total RNA was isolated from an independent infection experiment with three biological replicates using the Spectrum™ Plant Total RNA Kit (Sigma-Aldrich). RNA was DNaseI treated prior to cDNA synthesis to remove any potential genomic DNA contamination (Turbo DNA-free kit, Thermo Fisher Scientific). Relative quantification/Real-Time PCR (q-PCR) primers were designed for five reference genes and eight target genes using the Geneious software package (v10.0.3) (https://www.geneious.com) based on an annealing temperature of 60°C and short (< 120 bp) amplicons (supplemental file 13). As the *Psa* transcript content is low in a background of a large amount of total plant RNA, gene-specific reverse transcription (RT) primers were designed to prime the 1^st^ strand cDNA of the *Psa* specific transcripts [67]. Each RT primer (including reference genes – see supplemental file 13) was mixed and used as a cocktail for 1^st^ strand cDNA synthesis at a final concentration of 200 nM with 1 μg of total RNA in a 20 μl reaction according to manufacturing instructions (Superscript IV reverse transcription kit, Invitrogen). After heat denaturing of the RNA at 65°C for 5 min, RT primers were annealed at 55°C for 2 minutes and 1^st^ strand cDNA was synthesized at 53°C for 20 minutes followed by 15x dilution prior to qPCR. No RNase H step was included.

RT-qPCR was performed (four technical replicates per sample) using a LightCycler^®^ 480 Real-Time PCR System (Roche Diagnostics, Indianapolis, USA) using the LightCycler^®^ 480 SYBR Green I Master mix and amplified according to manufacturer’s instructions (60°C annealing for 10 s followed by extension at 72 °C for 20 s for 40 cycles) [68]. None of the water (negative template) controls or plant derived samples without *Psa* infection showed any amplification with cross points (CP) <35. Any samples with CP >35 were considered not expressed. Primer efficiencies were determined by serial template dilution. The best reference five genes were selected based on the RNA-seq experiment that showed low coefficient of variation (standard deviation divided by mean) and after RT-qPCR were validated using the Bestkeeper [69] and geNorm [70] tools. Based on this analysis, the geometric average CP of three genes was chosen (IYO_010670, IYO_009010 and IYO_002170) to represent the reference. For visual comparison between RT-qPCR and RPKM data, values were normalized by representing maximum expression as 100. Pearson correlation coefficients (r) between RT-qPCR and RPKM data were calculated in Microsoft Excel (CORREL function).

**Abbreviations**
AAD: amino acid adenylation protein
CEL: Conserved effector locus
CFU: Colony-forming units
EEL: Exchangeable effector locus
GABA: *γ*-aminobutyric acid
GMD: GDP-mannose dehydrogenase
HPCP: Hierarchical clustering on principal components
HPI: Hours post inoculation
EPS: Extracellular polysaccharide substances
IV: *in vitro*
LB: Lysogeny broth
LCA: Last common ancestor
NRP: Non-ribosomal peptide
PCA: Principal component analysis
*Psa*: *Pseudomonas syringae* pv. *actinidiae*
*Pss*: *Pseudomonas syringae* pv. *syringae*
Pto: *Pseudomonas syringae* pv. *tomato*
RPKM: Reads per kilobase per million
RT: Reverse transcription
RT-qPCR: Reverse transcription quantitative PCR
SNP: Single nucleotide polymorphism
T2SS: Type II secretion system
T2SP: Type II secreted proteins
T3SS: Type III Secretion System
T3SE: Type III Secreted Effectors
T6SS: Type VI Secretion System
TraDIS: Transposon directed insertion-site sequencing
VST: variance stabilizing transformation

## Ethics approval and consent to participate

All experiments using *Psa* were carried out with the permission of the Ministry for Primary Industries, New Zealand (CTO approval 12-05-17) and The Environmental Protection Authority, New Zealand (APP202231).

## Consent to publish

N/A

## Availability of data and materials

The RNA-seq experiment is described in BioProject PRJNA472664 with separate BioSamples for each time-point (SAMN09240241-97), reads can be downloaded from the Sequence Read Archive SRP148711 [71].

## Author contributions

BC, ACA, NN and CGP conceived of and designed the experiments. BC, RH-K, OvdL, XC, NN, ES and JJ carried out the experiments and generated the data. PM, LB, NN, EHR and MDT analyzed the data. MDT, PM, NN, SN, EHR, and ACA wrote the paper.

## Acknowledgements

We would like to thank Dr Joanna K. Bowen and Amali Thrimawithana (Plant and Food Research) for critically reviewing this manuscript.

---

**Additional file 1:**
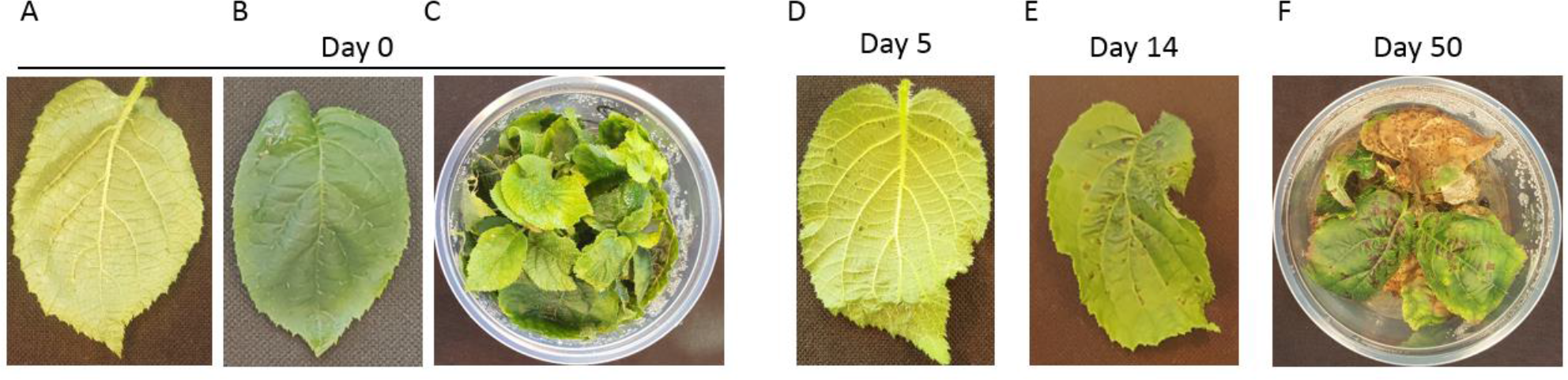
Images illustrating the time course of symptom development of kiwifruit plantlets infected with *Psa*. (A) Abaxial side of leaf at day 0; (B) Adaxial side of leaf at day (0); (C) Pottle containing plantlets at day 0; (D) Abaxial side of leaf five days post inoculation (DPI) with water soaked lesions appearing; (E) Adaxial side of leaf 14 DPI with necrotic lesions present; (F) Plantlets 50 DPI (pptx).

**Additional file 2:** Reads per Kilobase per Million (RPKM) for all *Psa* genes (sheet 1) and k-means analysis for those genes with >50 RPKM for at least one time point (sheet 2) (xlsx).

**Additional file 3:**
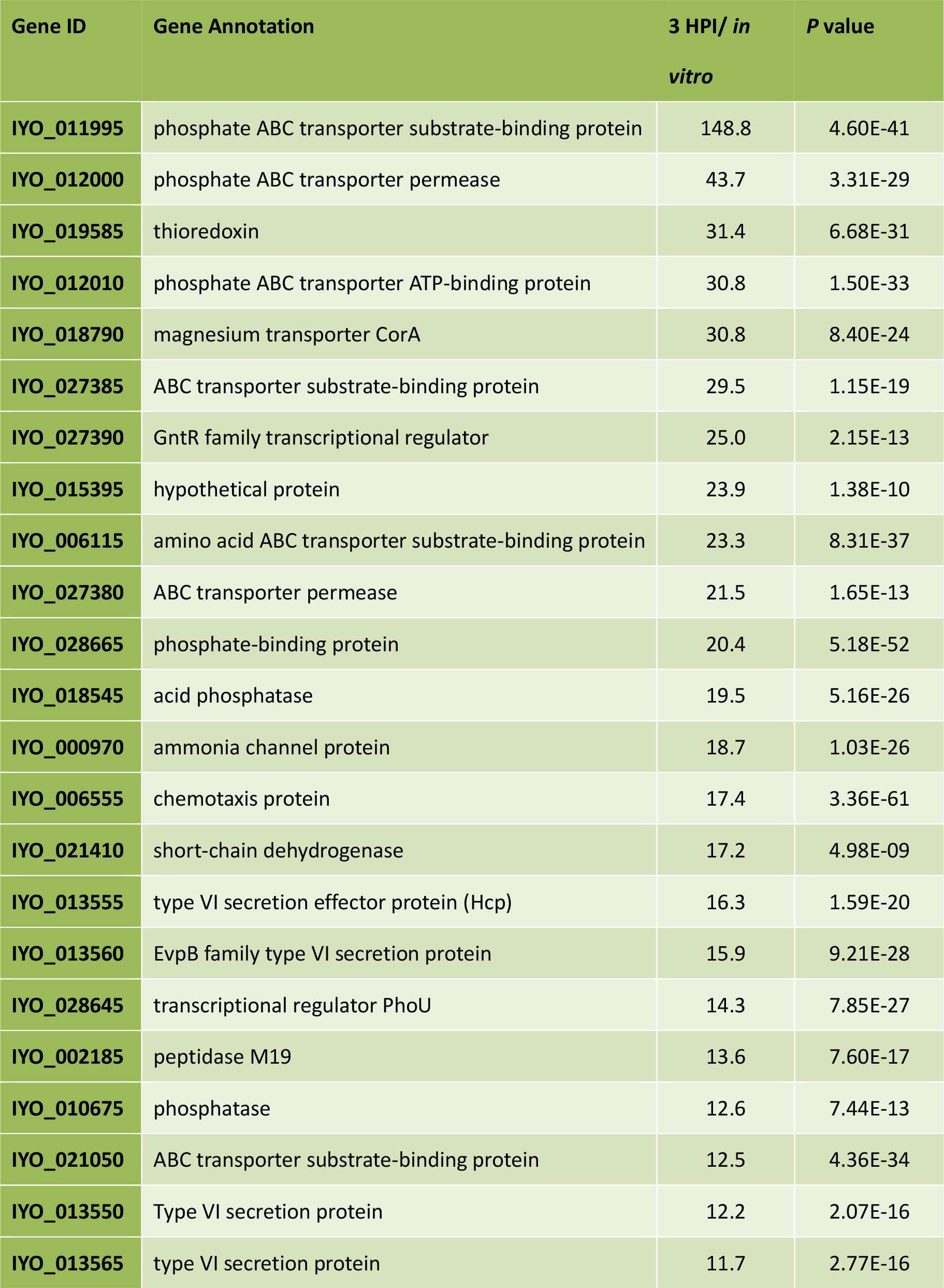
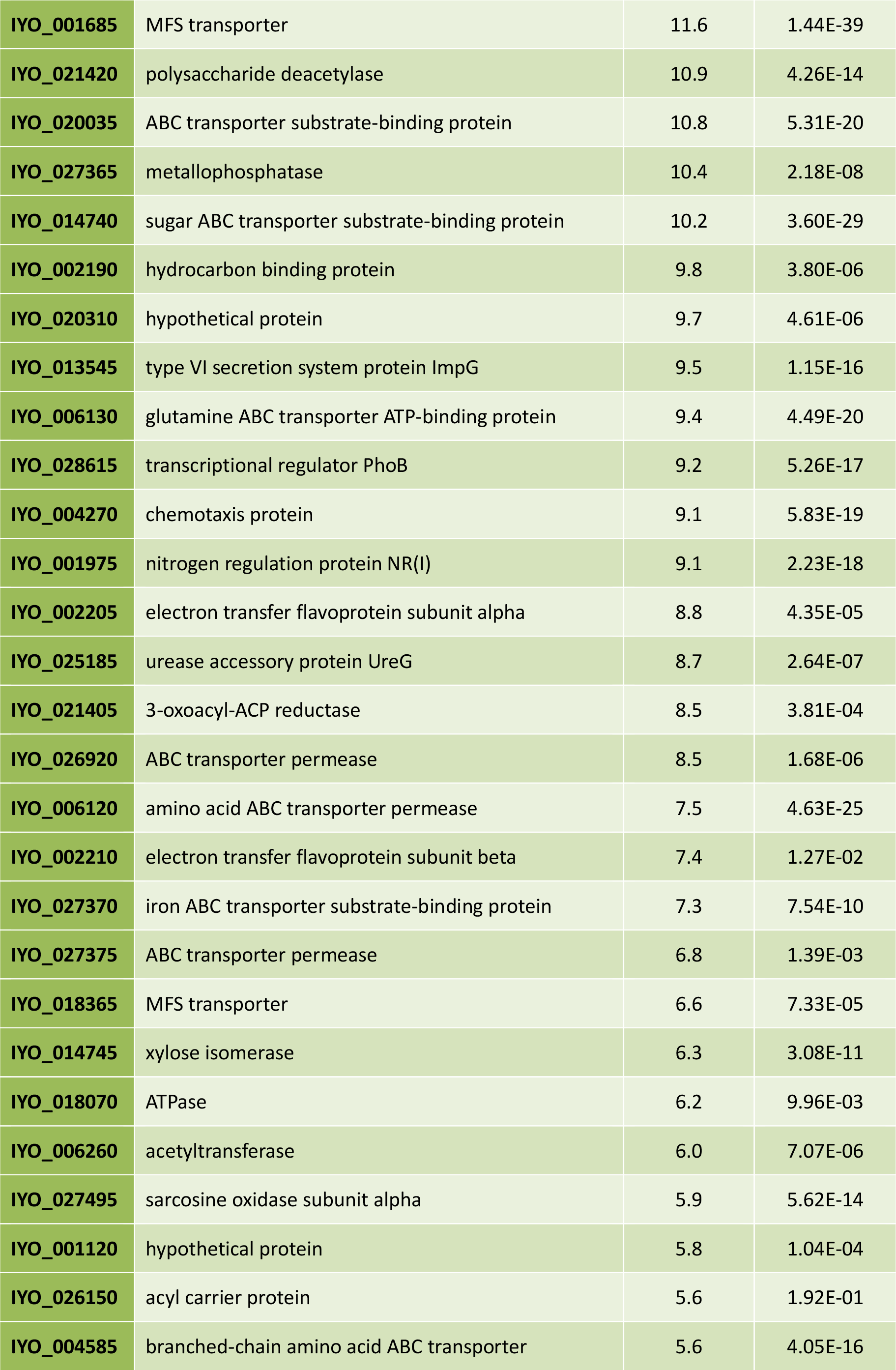
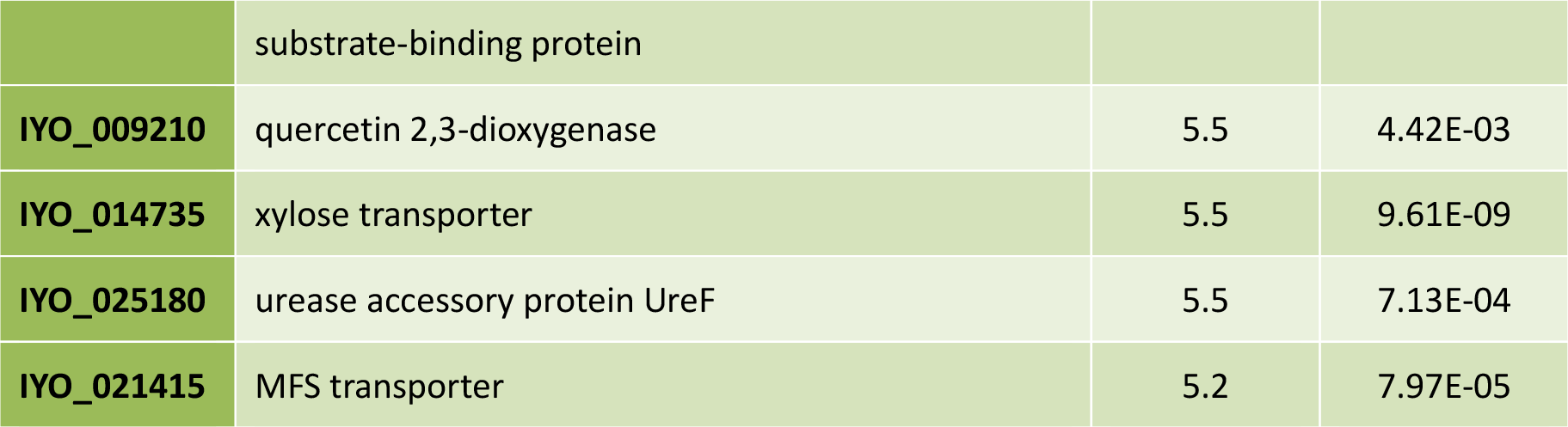
Early induced genes ranked by the ratio of expression at three hours post infection (HPI) compared with *in vitro* (cutoff 5-fold) (docx).

**Additional file 4:**
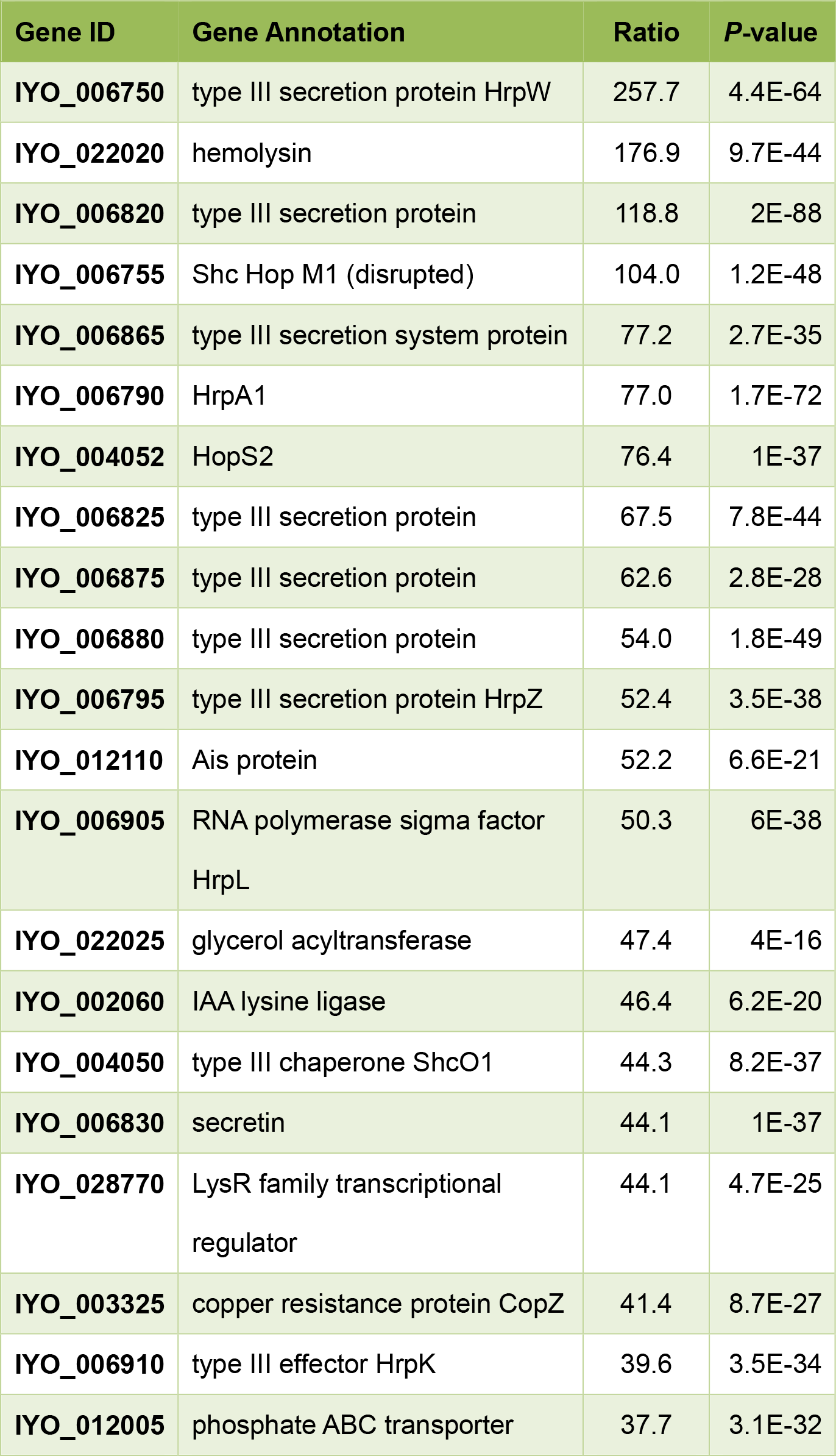
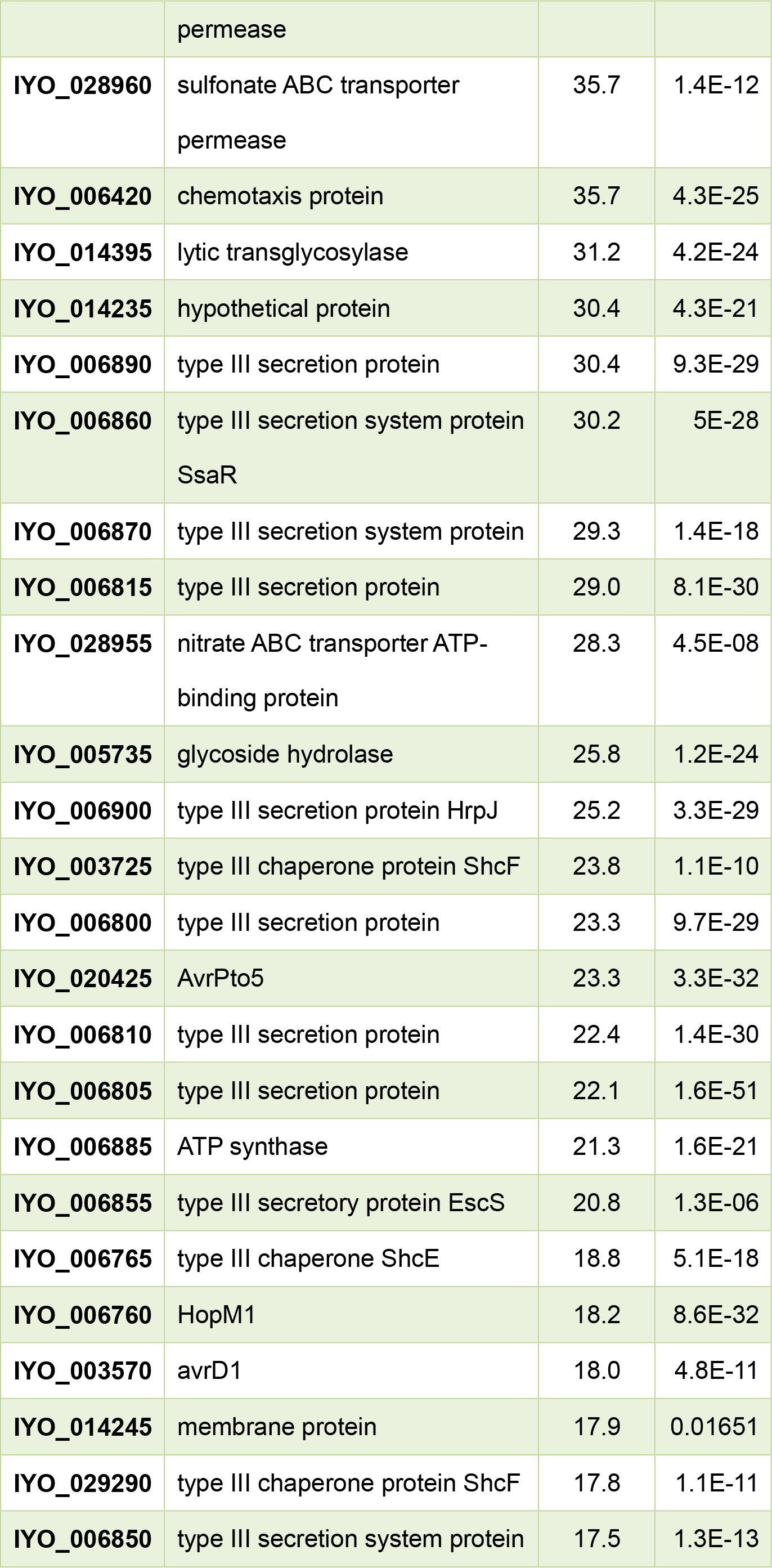
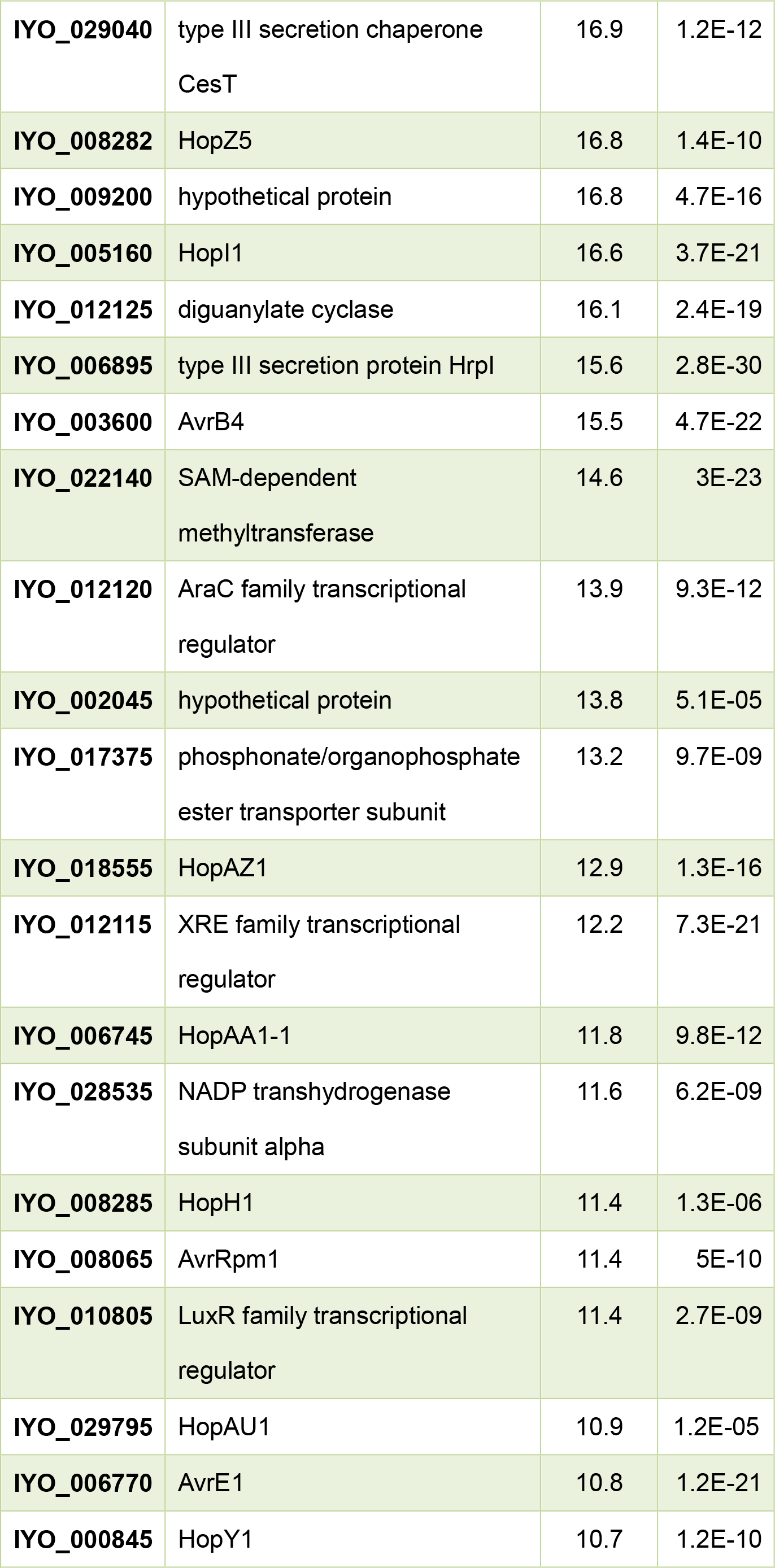
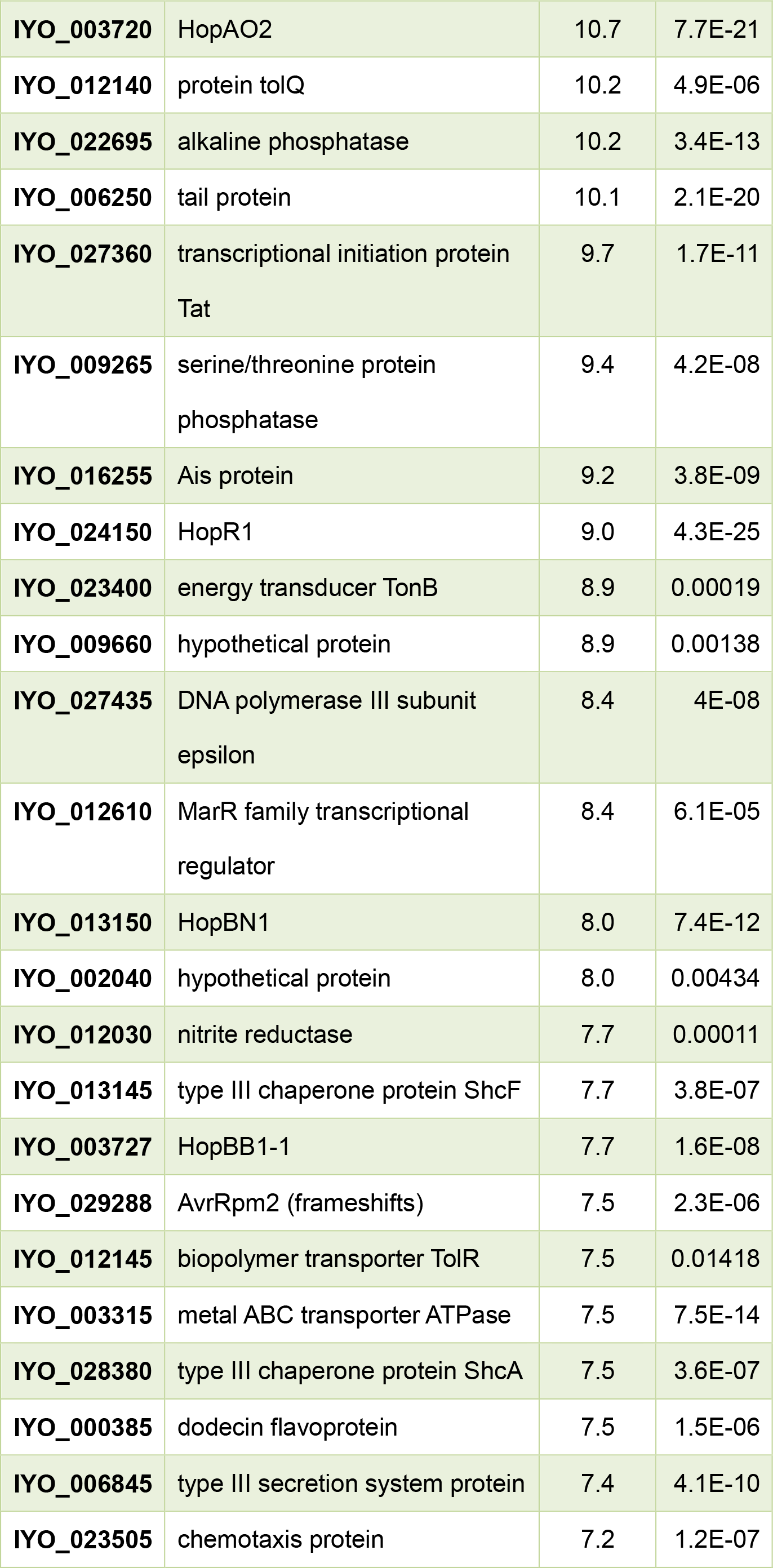
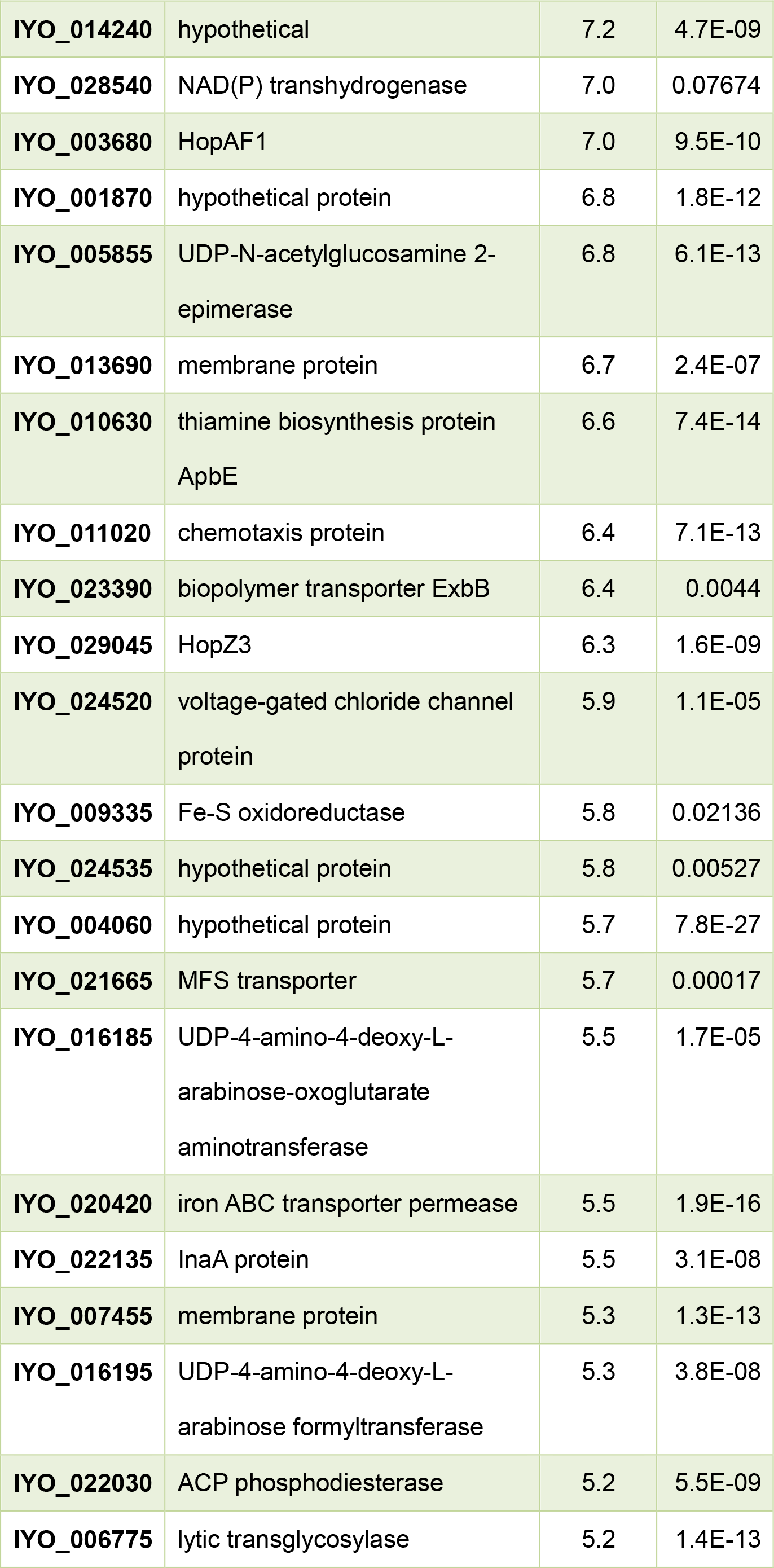
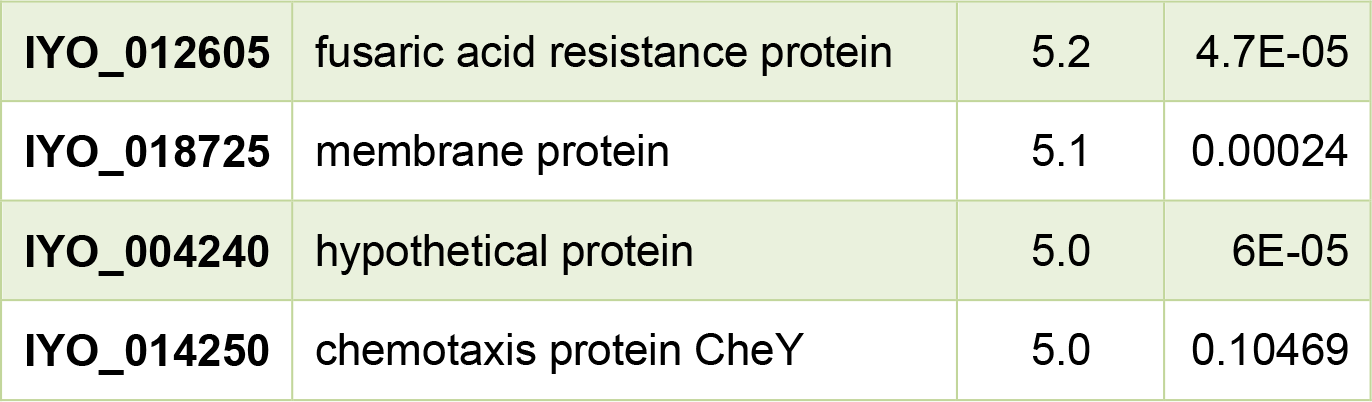
Genes most highly upregulated in the mid phase of the infection time course (3-24 hours post infection, (HPI)). Genes are ranked on the ratio of maximum Reads Per Kilobase per Million over that time period compared with *in vitro* expression (cutoff 5-fold) (docx).

**Additional file 5:**
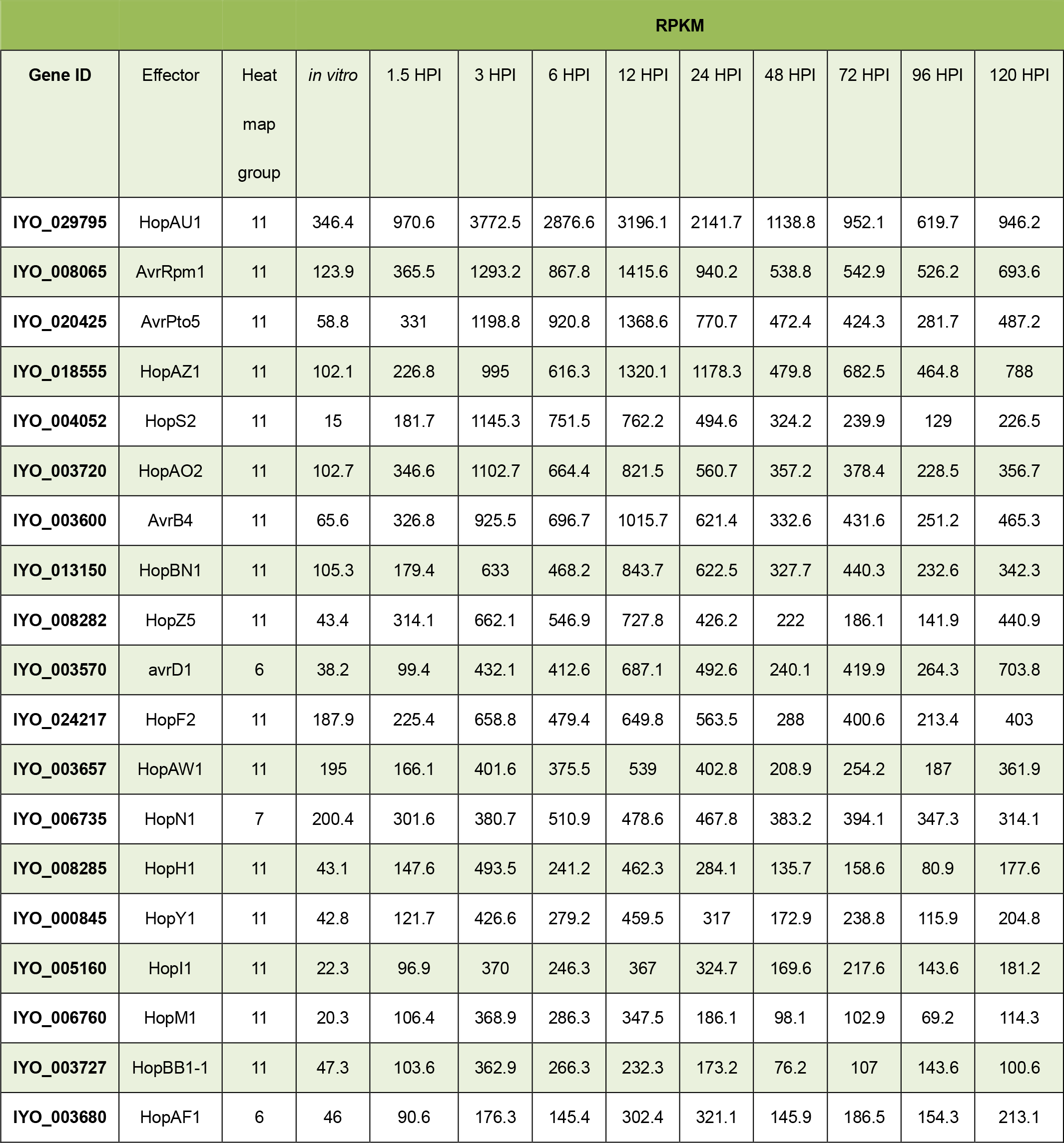
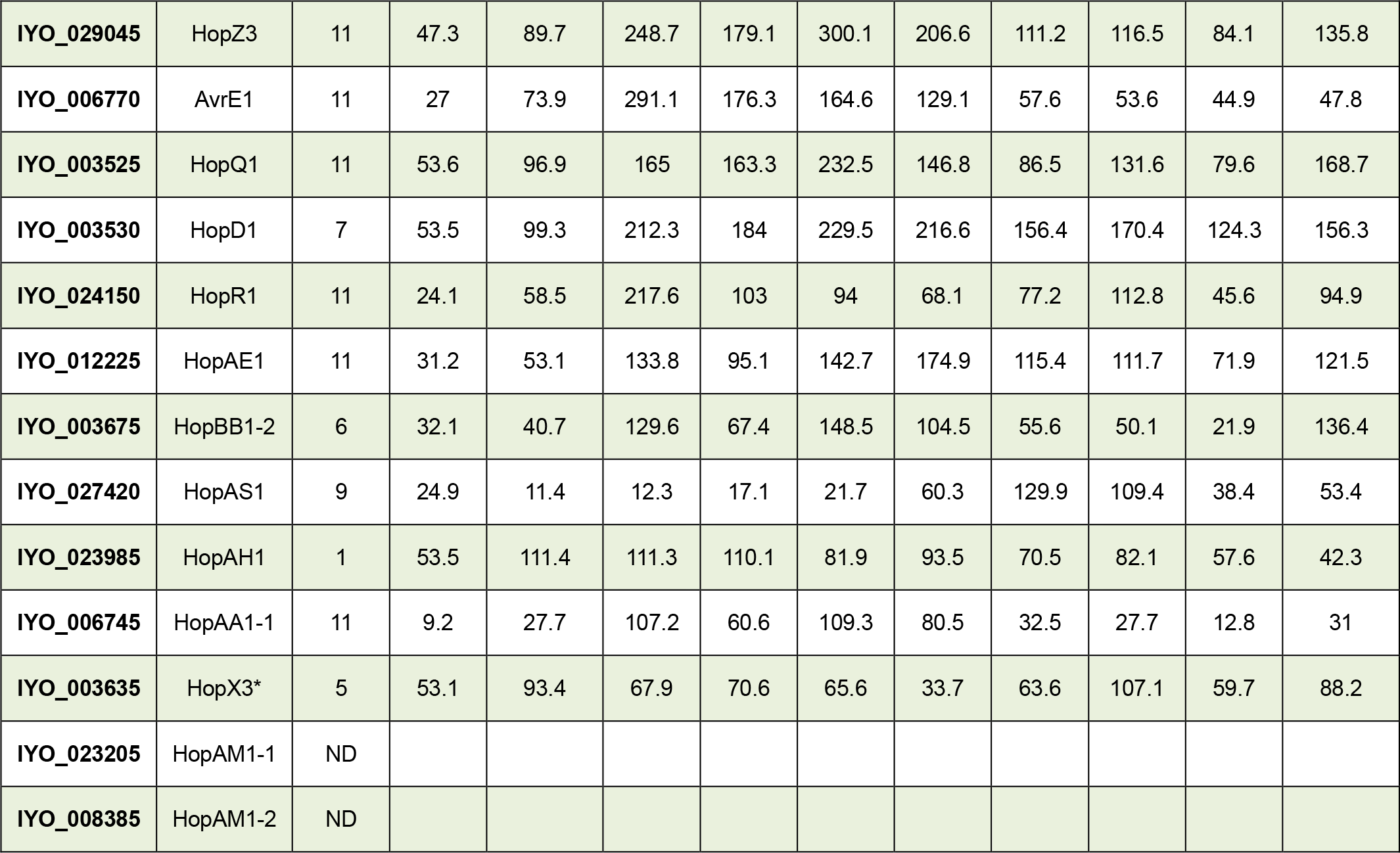
Expression levels of individual *Psa* effectors over the infection time course. Effectors are ranked by the highest level of expression between 3 and 12 hours post infection (HPI) in reads per kilobase per million (RPKM). Effectors likely to be disrupted or pseudogenes were not included (docx).

**Additional file 6:**
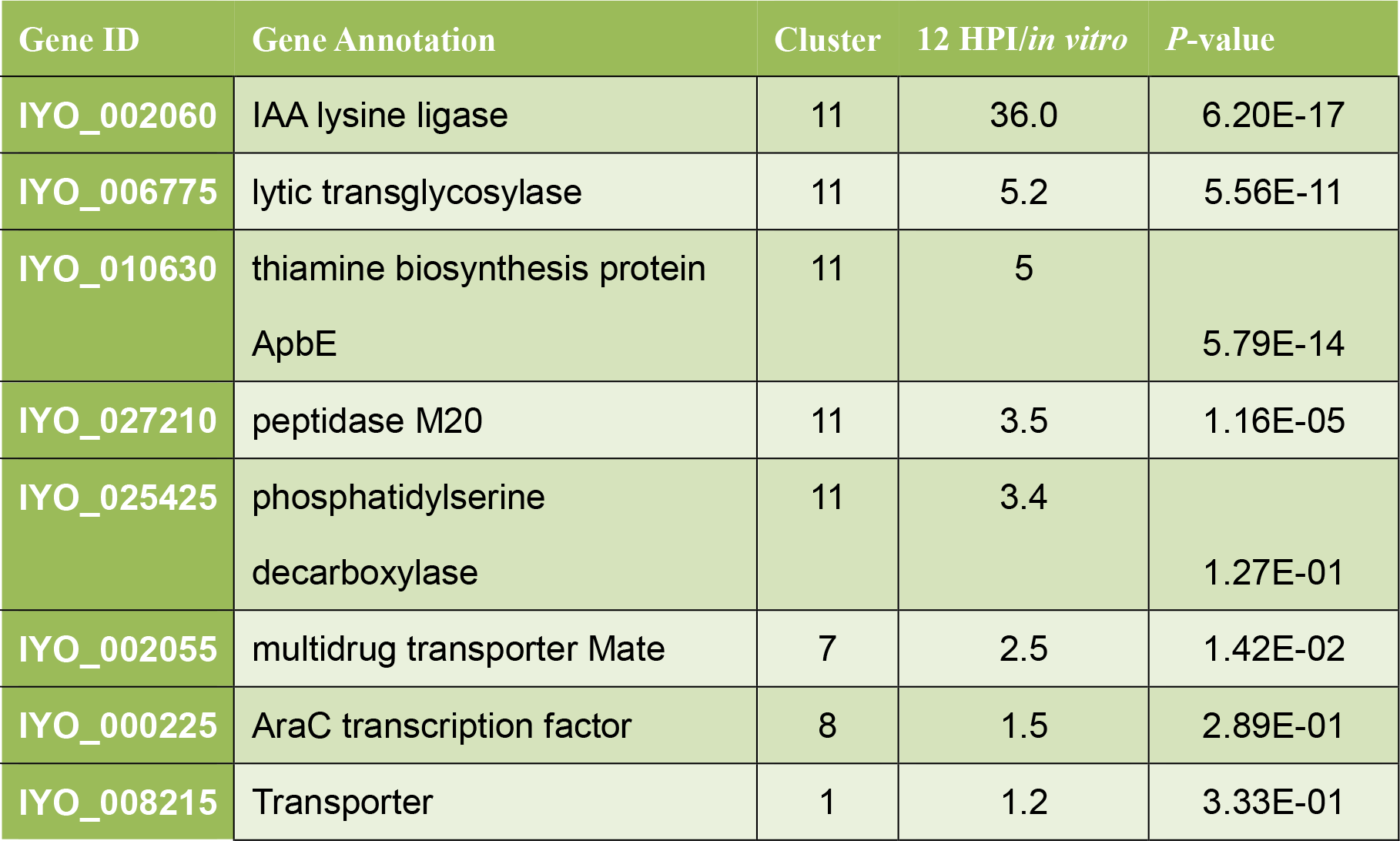
Expression of non-effector genes with upstream HrpL boxes. Genes are ranked by the ration of expression at 12 hours post infection (HPI) compared with *in vitro* expression (docx).

**Additional file 7:**
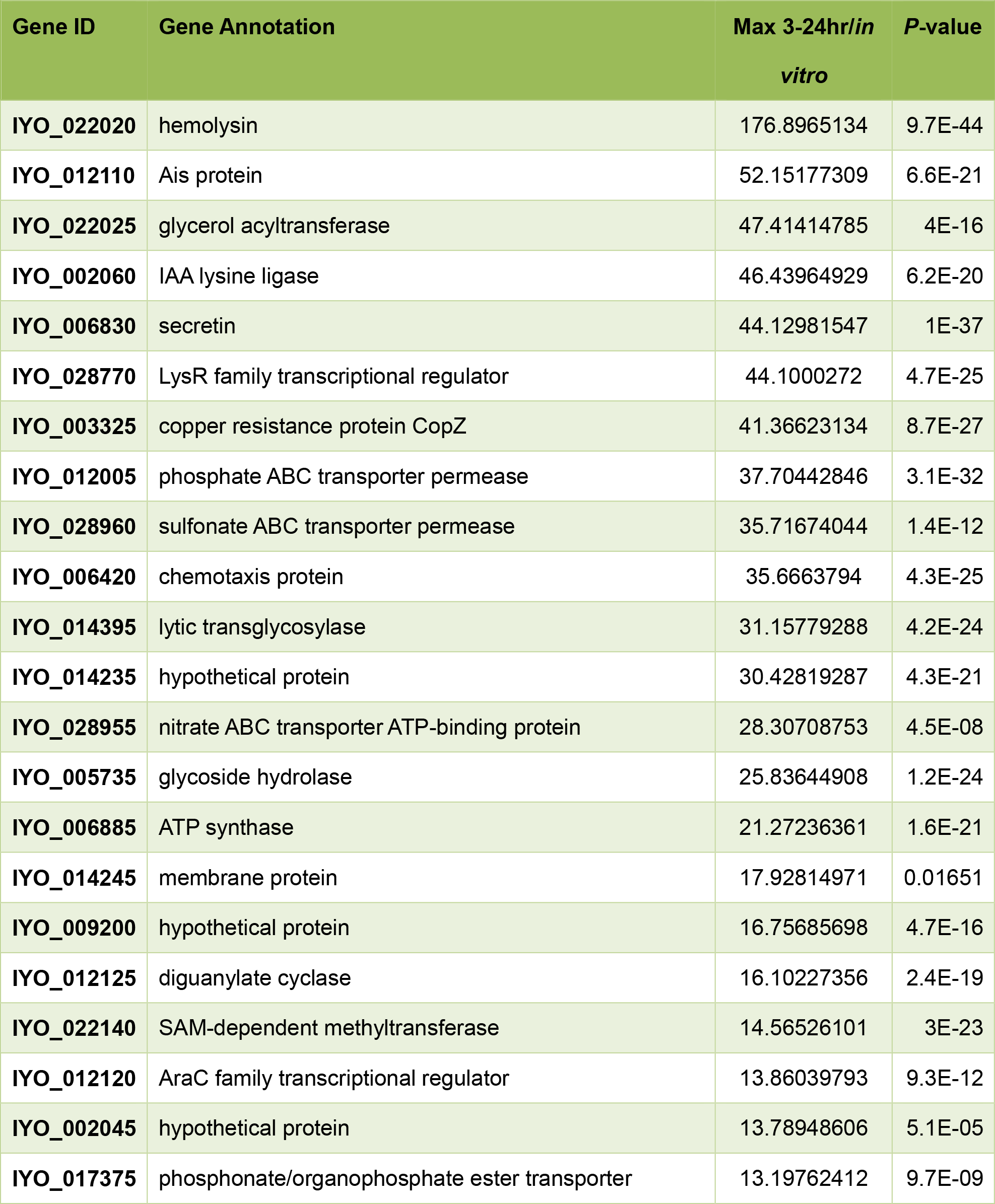
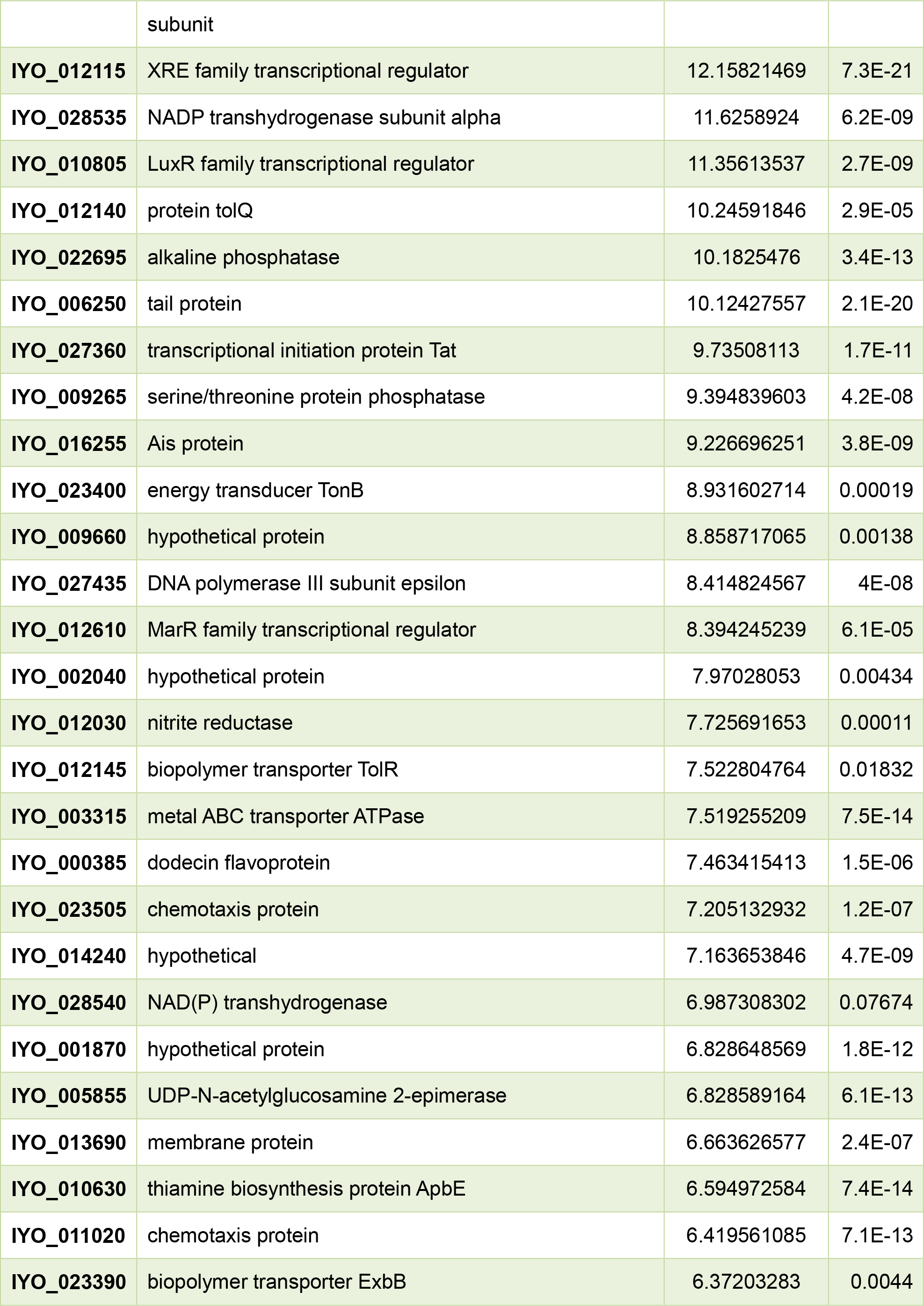
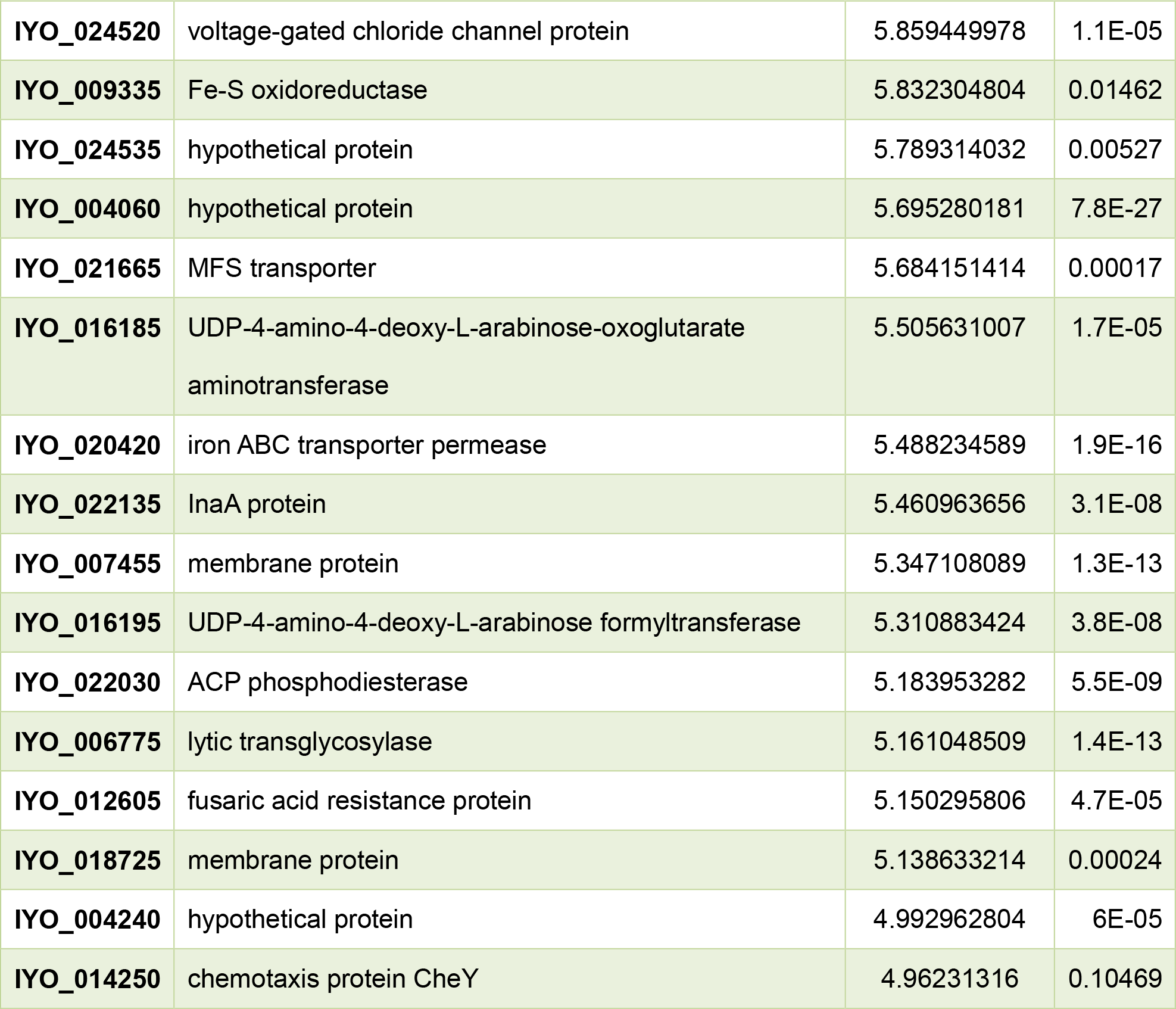
Genes expressed in the mid phase of infection that do not encode the T3SS or T3SEs. Effectors were ranked by the highest level of expression between 3 and 24 HPI compared with *in vitro* (cutoff 5-fold) (docx).

**Additional file 8:**
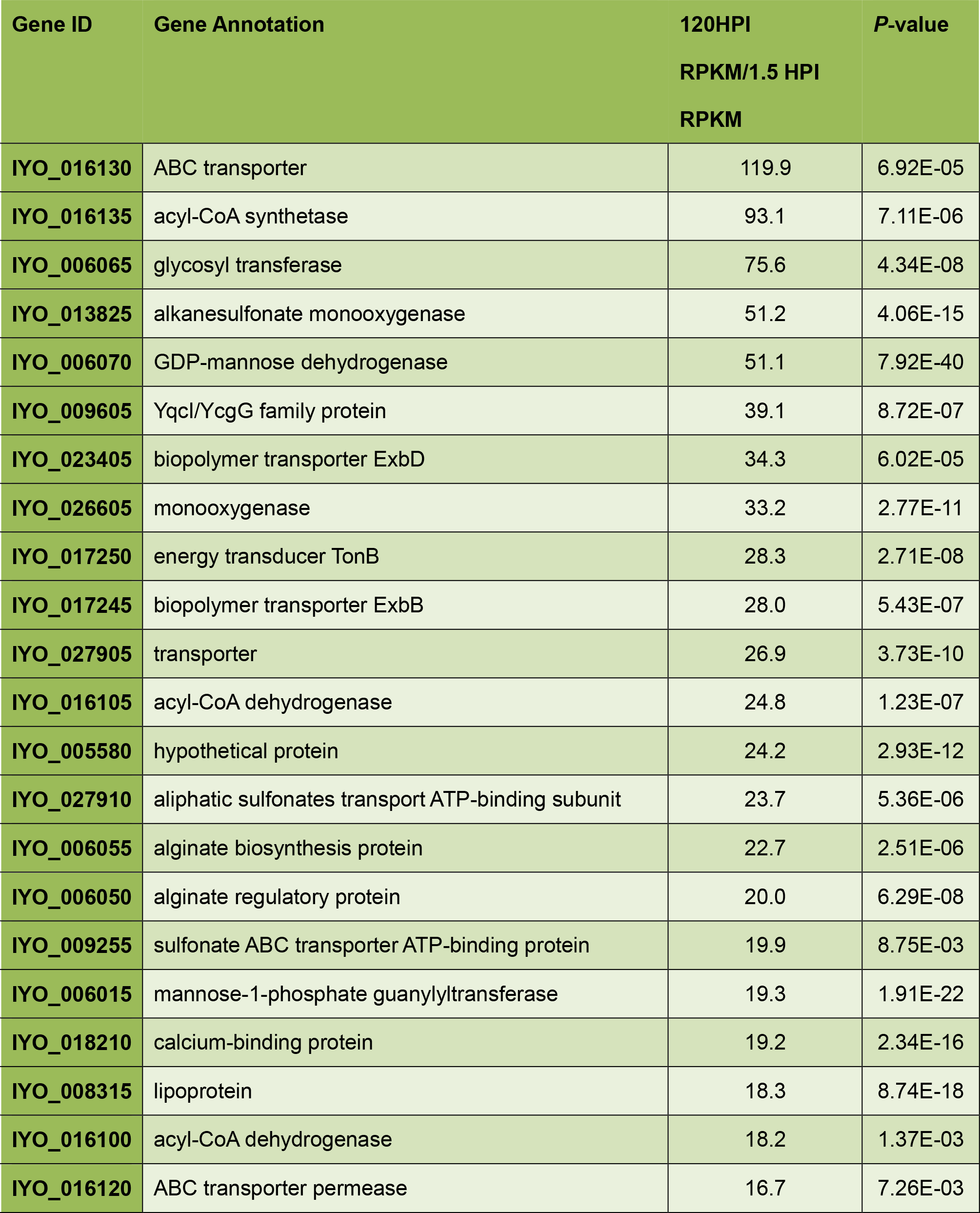
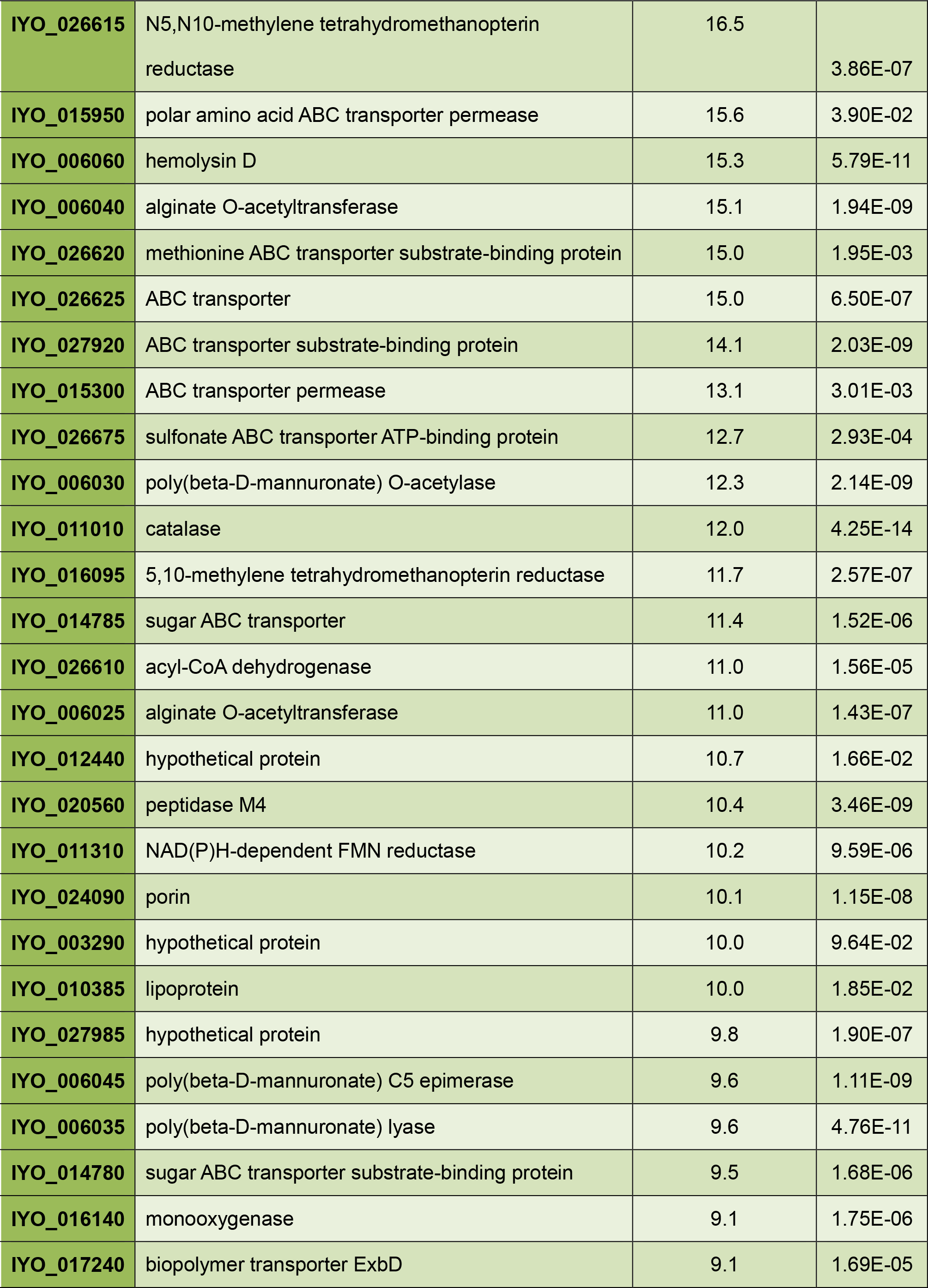
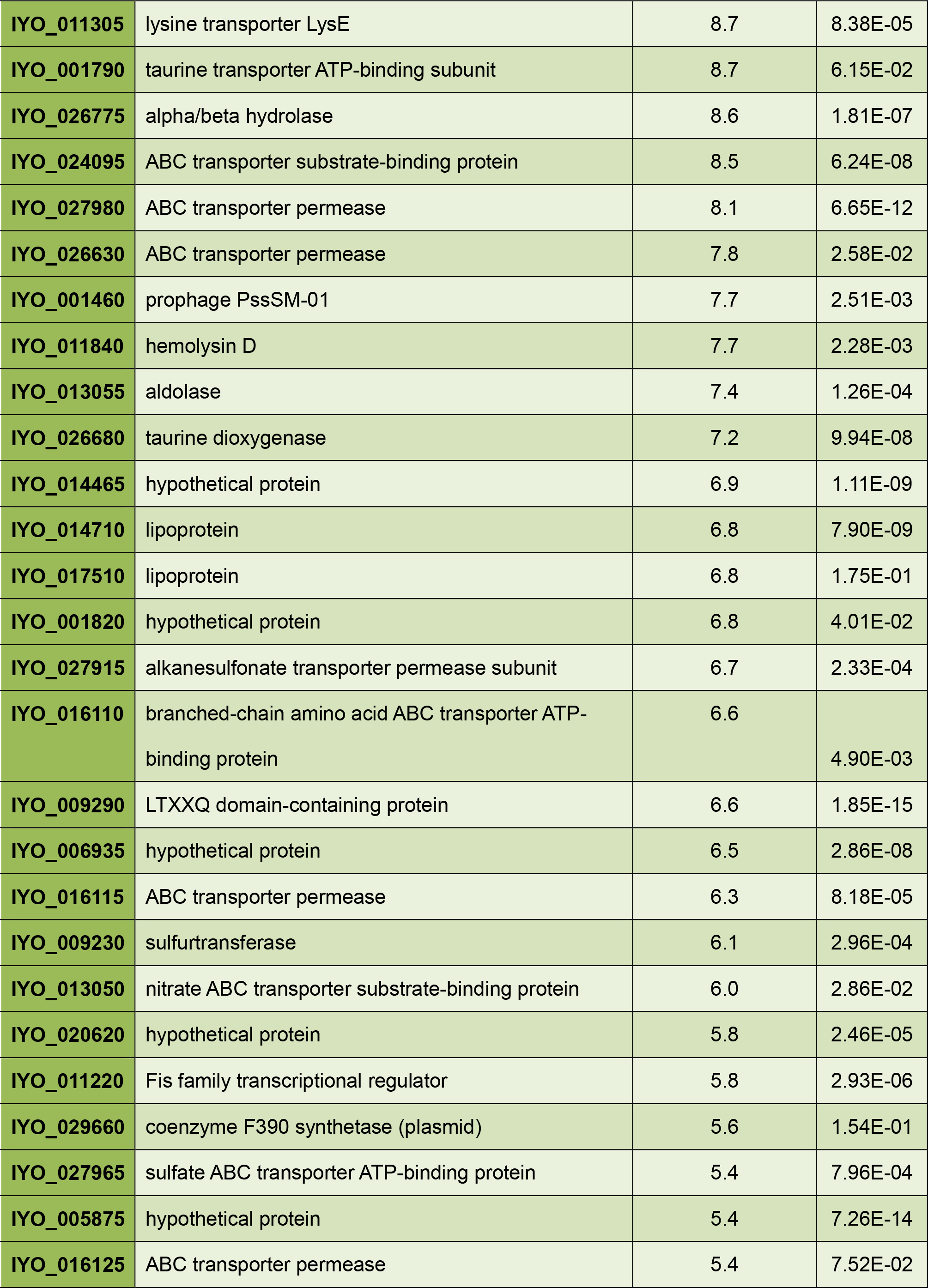
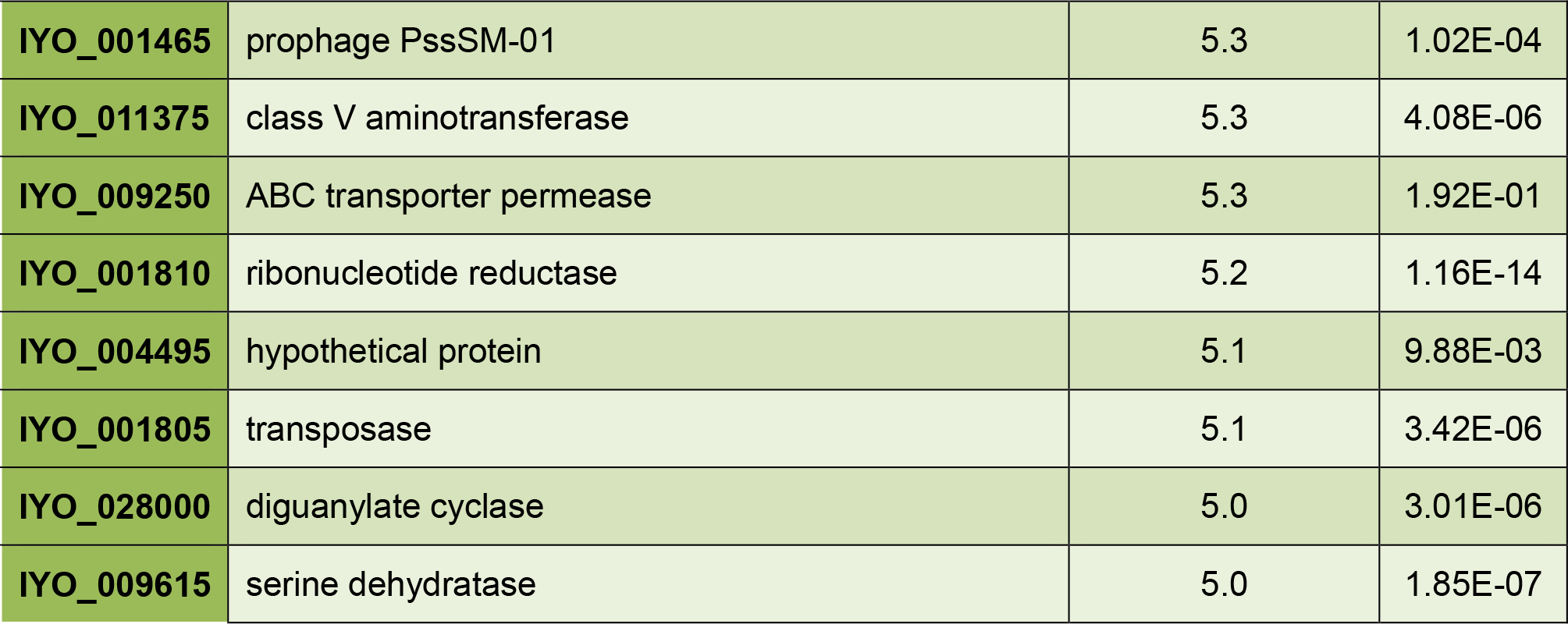
Genes upregulated late in the time course. Genes were ranked based on the ratio of expression at 120 hours post infection (HPI) compared with 1.5 HPI (docx).

**Additional file 9:**
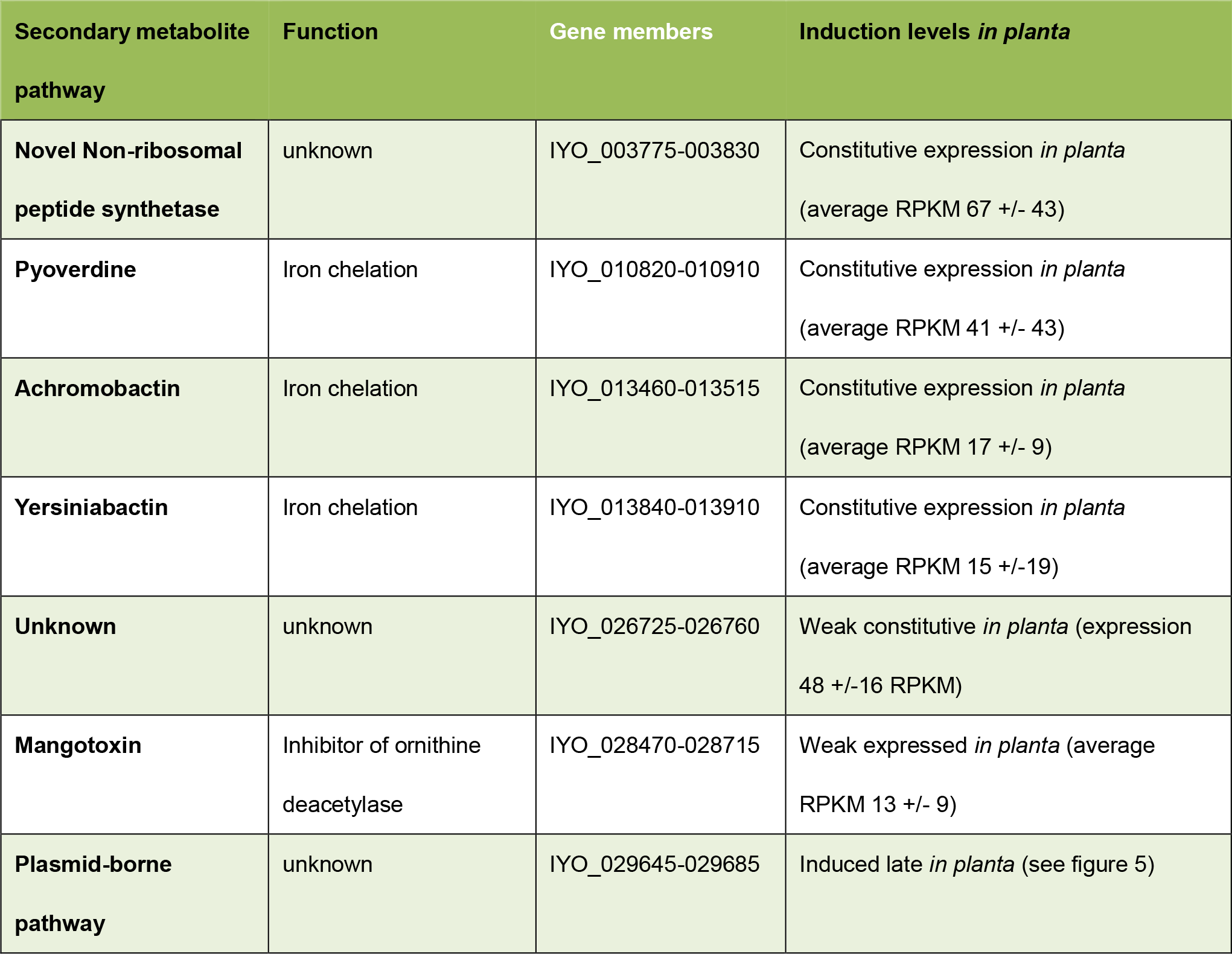
Expression levels of secondary metabolite gene clusters. Means of reads per kilobase per million (RPKM) for each gene across all time points with standard deviations (docx).

**Additional file 10:**
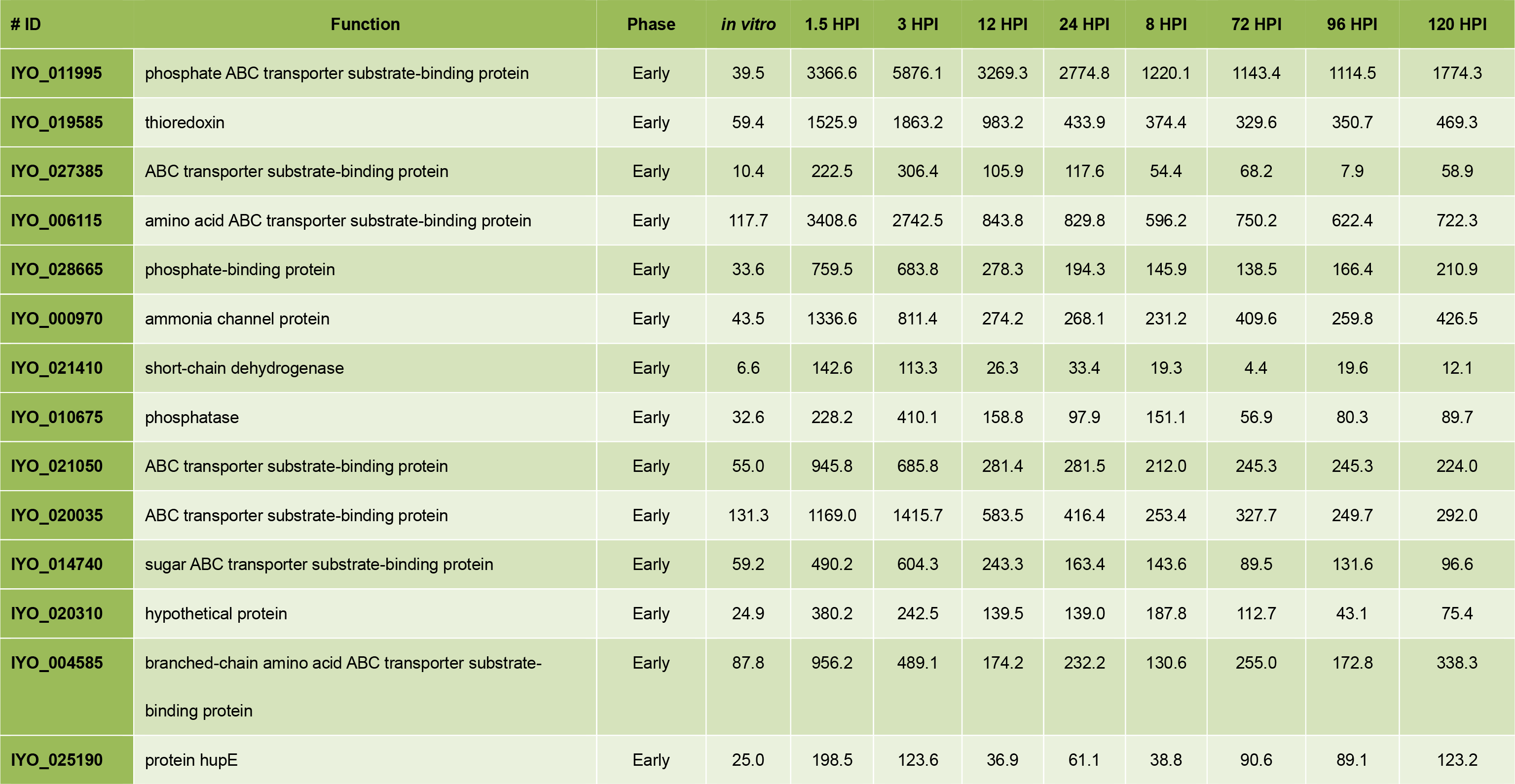
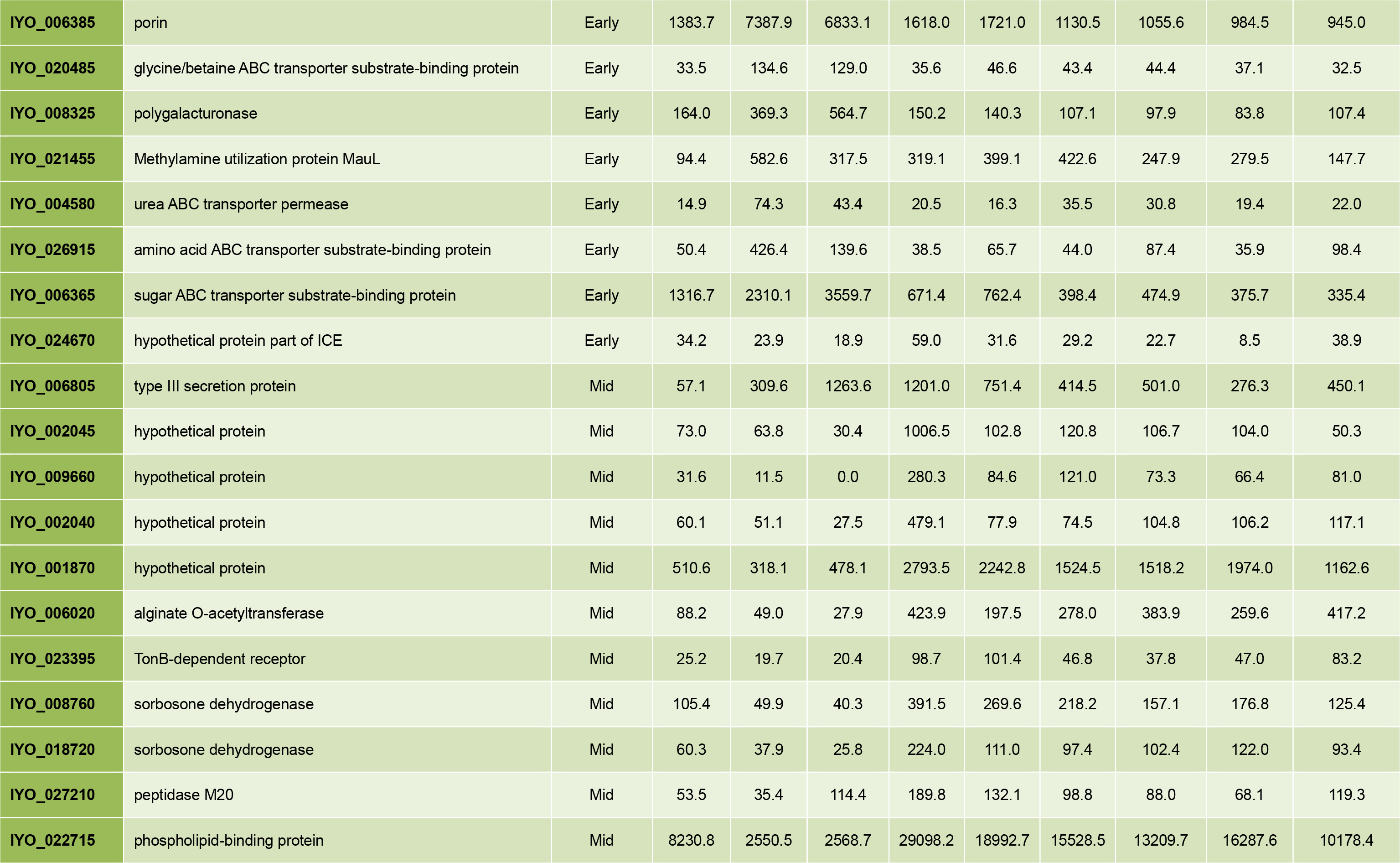
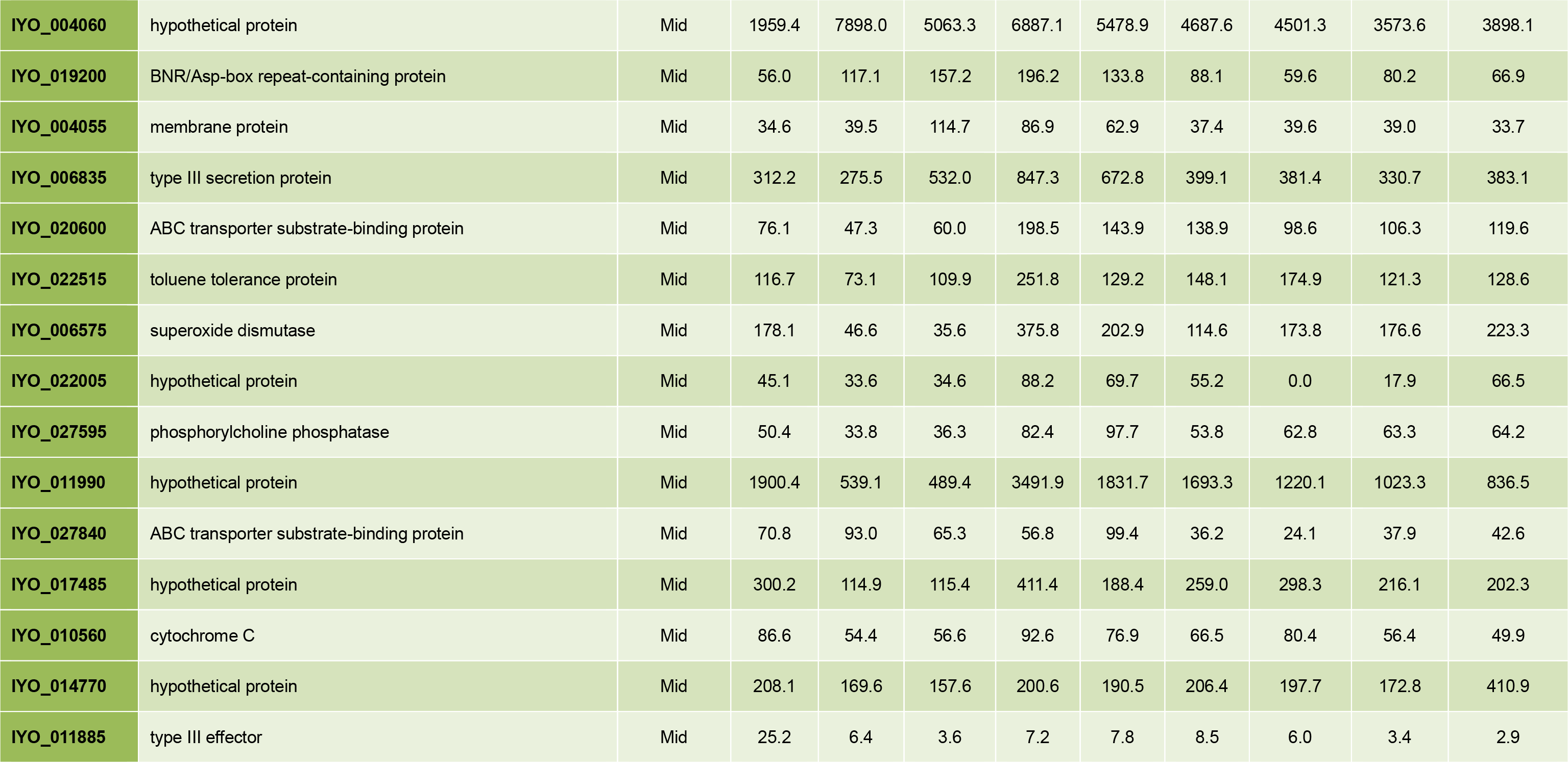
Reads per kilobase per million (RPKM) values of genes encoding proteins predicted to be secreted via T2SS (docx).

**Additional file 11:**
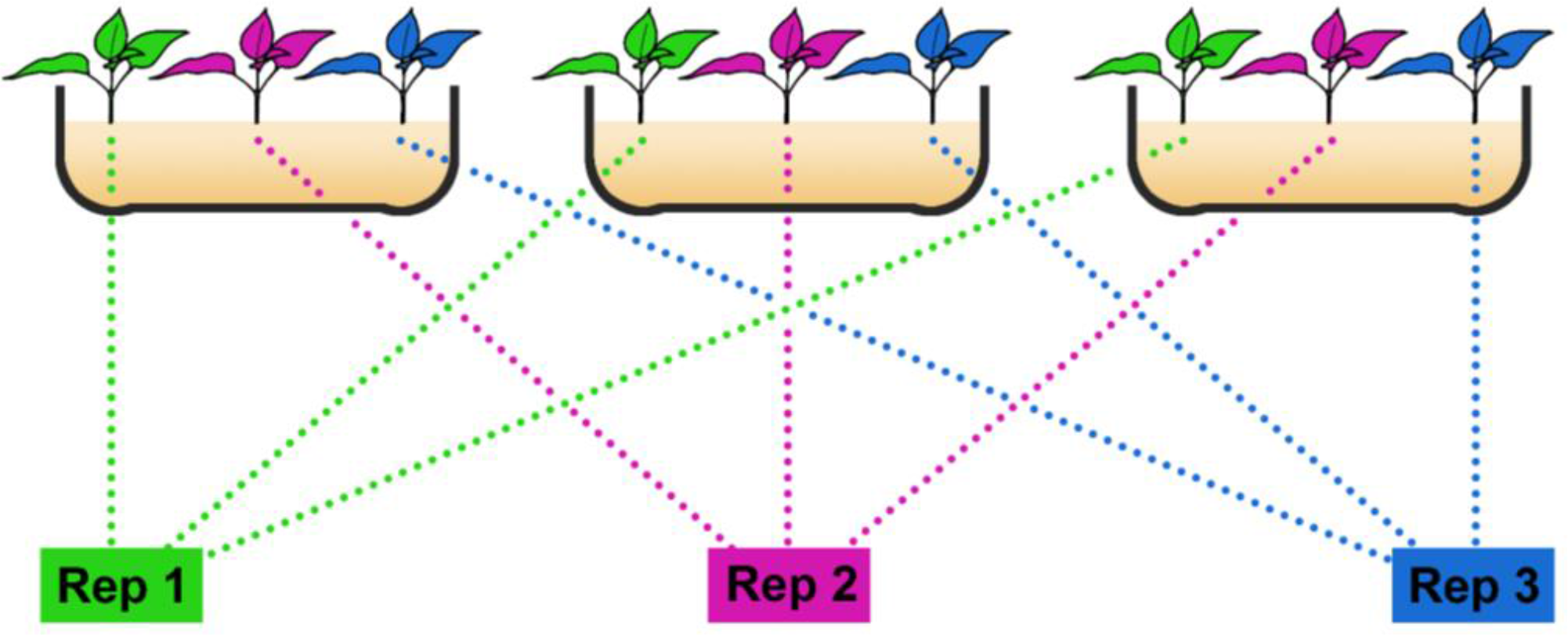
RNA-seq experimental design. Three pottles, each with three plantlets, were inoculated for each time point. For RNA extraction, one plantlet from each pottle, was harvested and combined for each of three biological replicates (docx).

**Additional file 12:**
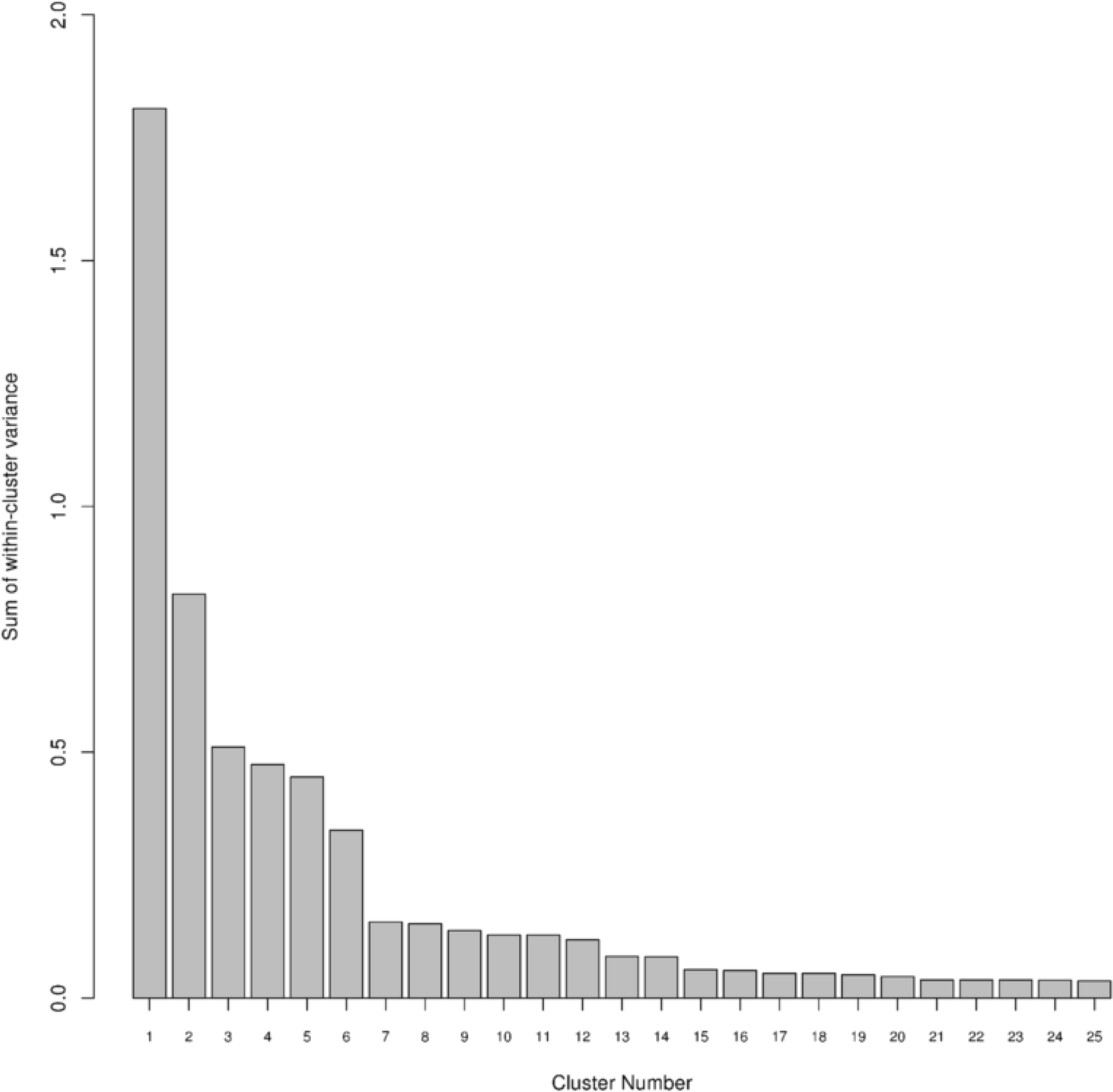
A bar plot of the inertia gain using the sum of the within-group variance with increasing cluster number (x-axis) produced using Hierarchical clustering on principal components (tiff).

**Additional file 13:** A spreadsheet containing all the primers used for reverse transcription and reverse transcription quantitative PCR. The data includes the gene chosen, locus ID number, primer sequences, product size and PCR efficiency (xlsx).

